# DNA supercoiling links transcription and chromatin architecture during human stem cell differentiation

**DOI:** 10.64898/2026.02.12.705511

**Authors:** Consuelo Perez, Pierre Murat, Andrew Zeller, Kim C. Liu, Alastair Crisp, Julian E. Sale

**Affiliations:** Division of Protein & Nucleic Acid Chemistry, Medical Research Council Laboratory of Molecular Biology, Francis Crick Avenue, Cambridge, CB2 0QH, UK

**Author notes:** Current address: Wellcome Sanger Institute, Wellcome Trust Genome Campus, Hinxton, CB10 1RQ, UK. Joint first authors (Consuelo Perez, Pierre Murat).

## Abstract

Transcription imposes torsional stress on chromatin, leading to over– or under-winding of the DNA helix. Yet, how supercoiling evolves during dynamic changes in gene expression, and how it influences three-dimensional chromatin contacts in the densely packed human genome, remains unclear. Here, we map genome-wide negative supercoiling and chromatin interactions at high resolution in human embryonic stem cells, using the transcriptional changes that drive differentiation to explore their interplay at multiple scales. We demonstrate that gene activation increases negative supercoiling despite enhanced topoisomerase recruitment. Moreover, elevated negative supercoiling correlates with increased DNA contacts within domains at multiple scales, from genes to topologically associating domains, consistent with transcription-generated torsional stress enhancing contact frequency. Thus, negative supercoiling likely contributes to the genome architectural remodelling that accompanies the execution of developmental programs.

## Introduction

Molecular machines that translocate on DNA, such as RNA or DNA polymerase, generate torsional forces along the axis of the helix. In a short linear section of DNA this torque will be dissipated by free rotation of the ends of the molecule. However, when rotation of the helix is constrained, torsional stress accumulates leading to over– or under-winding of the helix, known as supercoiling. Thus, in constrained transcribed domains, positive supercoiling (overwinding) is expected to accumulate ahead of RNA polymerase II (RNAPII) and negative supercoiling (underwinding) behind, as described by the twin domain model (Figure 1A) (Liu and Wang, 1987; Janissen *et al*., 2024; Gilbert and Marenduzzo, 2025).

**Figure 1.**
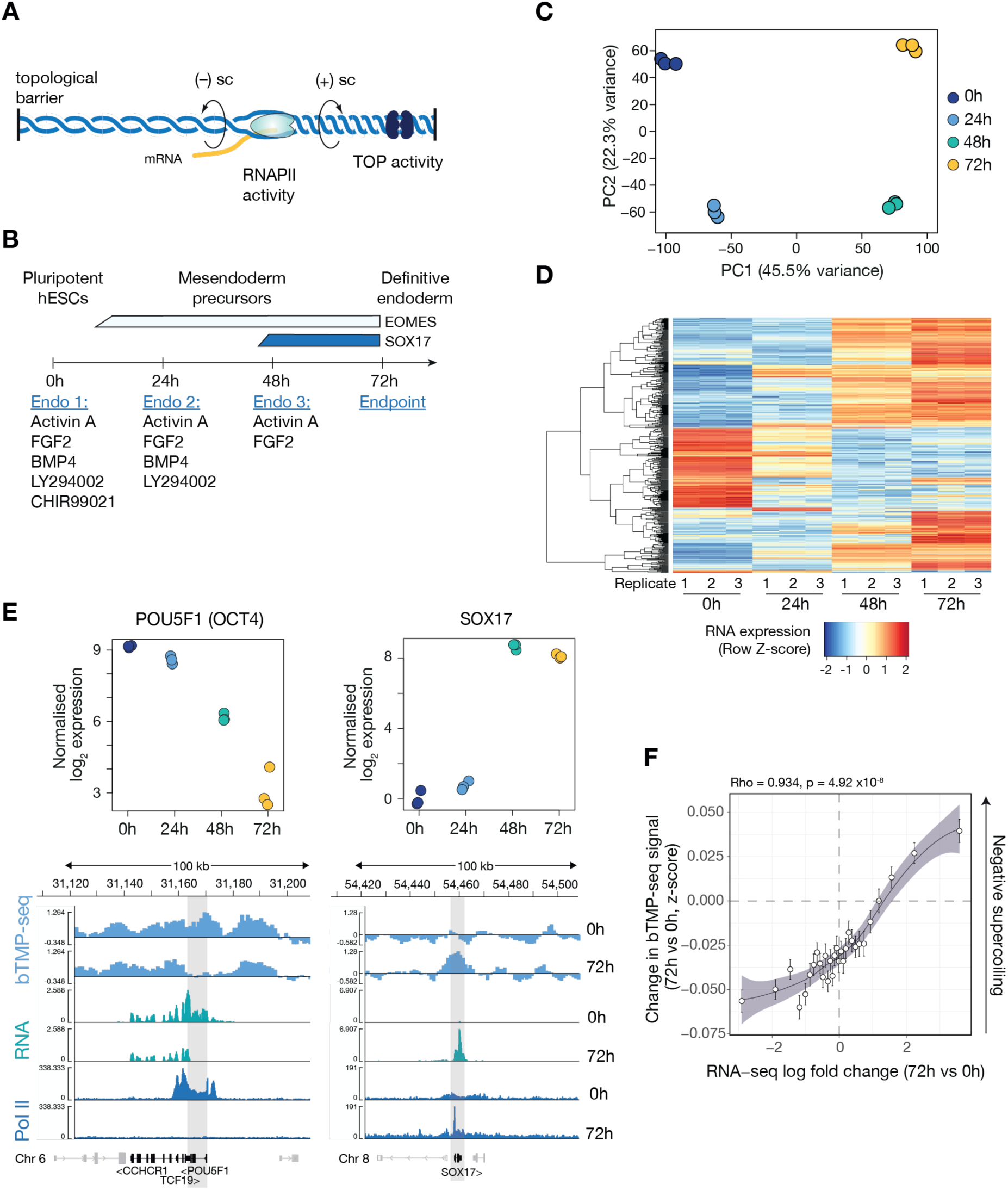
Monitoring DNA supercoiling in hESCs differentiating to definitive endoderm. A. Scheme summarising the twin domain model: movement of RNAPII creates positive supercoiling in front and negative supercoiling behind the elongating polymerase that is counteracted by topoisomerase activity. B. Schematic of the timeline and medium composition changes used to generate definitive endoderm from H9 human embryonic stem cells. The bars above the timeline indicate expected changes in expression of key transcription factors. Adapted from Eldridge et al., 2020 (Eldridge *et al*., 2020). C. Principal component analysis of RNA-seq from H9 cells differentiating to endoderm. Each dot represents an independent biological replicate collected at the indicated time point. D. Heatmap and unsupervised clustering of the top 500 most differentially expressed genes through H9 endoderm differentiation. E. Example changes in bTMP-seq signal around the *POU5F1* and *SOX17* loci as a function of differentiation stage. The upper panels show the expression level of *POU5F1* and *SOX17* in samples collected every 24h across endoderm differentiation. Each point represents the normalised log_2_ RNA-seq expression values for one of three biological replicates at each time point. Lower panels show Integrative Genomics Viewer (IGV) browser tracks of the normalised supercoiling score (light blue), RNA-seq (green), and RNAPII (blue) around the *SOX17* and *POU5F1* loci (grey zones) in pluripotent (0h) and hESCs differentiated to definitive endoderm (72h). UCSC genes are marked with black squares (and non-coding RNAs in grey) at the bottom of the panel. Genome coordinates are shown at the top. F. Genome wide correlation between change in supercoiling score (ΔbTMP-seq signal, z-score) and log_2_ fold change in the RNA-seq signal in the body of long genes (≥30 kb) between the endoderm (72h) and undifferentiated (0h) states. Fit: LOESS; correlation: Spearman’s ρ.

Supercoiling is counteracted by topoisomerases, which relax positive or negative superhelicity by a transient strand cleavage and strand passage mechanism (Pommier *et al*., 2016, 2022). In low-output genes Topoisomerase I (Topo I) is sufficient to relieve torsional stress, whereas highly transcribed genes additionally recruit Topo II (Kouzine *et al*., 2013). Topo I physically associates with, and is stimulated by, the RNAPII complex (Baranello *et al*., 2016), directly coupling resolution of supercoiling with transcription. Nonetheless, evidence from an engineered system based on autonomously replicating episomes has shown that accumulation of supercoiling in a highly transcribed gene can outstrip topoisomerase recruitment (Kouzine *et al*., 2008). Further, mapping approaches exploiting the differential intercalation of psoralen into DNA as a function of its superhelicity (Sinden *et al*., 1980) indicated that the steady-state levels and distribution of supercoiling varies in bacterial (Visser *et al*., 2022; Fu *et al*., 2024), yeast (Bermúdez *et al*., 2010; Achar *et al*., 2020) and human genomes (Naughton *et al*., 2013; Yao *et al*., 2025), with accumulation of negative supercoiling being associated with transcriptionally active domains.

Energy stored as helical torsion of the DNA double helix can be converted into writhe, forming plectoneme structures, the position of which is modulated by sequence context (Kim *et al*., 2018). This has been proposed as a driver of increased physical contacts between loci within supercoiled domains (Benedetti *et al*., 2014). Chromatin contacts contribute to the regulation of gene expression both at short-range, within genes, and at long-range, bridging enhancers and promoters over distances that can extend to megabases (Lettice *et al*., 2003). Functional chromatin interactions are promoted by the compartmentalisation and three-dimensional organisation of the genome into large-scale structures (Lieberman-Aiden *et al*., 2009) known as topologically associating domains (TADs) (Dixon *et al*., 2012; Nora *et al*., 2012; Sexton *et al*., 2012), with the promoter and regulatory elements of a given gene generally found in the same TAD (Lupiáñez *et al*., 2015). TADs are nested within two computationally defined genomic compartments, A and B, which broadly correlate with transcriptionally active and heterochromatic regions of the genome respectively (Lieberman-Aiden *et al*., 2009). The arrangement of TADs at the megabase scale is strongly conserved across cell types suggesting that they arise as a fundamental property of the genome (Dixon *et al*., 2012; Nora *et al*., 2012), although sub-megabase rearrangements of TAD structure and cis-regulatory contacts occur during differentiation (Phillips-Cremins *et al*., 2013; Freire-Pritchett *et al*., 2017). The loop extrusion activity of cohesin, constrained by CTCF binding at convergently oriented motifs (Dixon *et al*., 2012; Vietri Rudan *et al*., 2015), provides a mechanistic framework for TAD and chromatin loop formation. However, current cohesin-based extrusion models do not fully explain the observed asymmetry in contact directionality, as cohesin is generally thought to extrude DNA bidirectionally until halted by CTCF. While this explains loop anchoring, the potential contribution of physiologically generated DNA supercoiling to loop formation and to both local and long-range genomic contacts has remained less clear (Benedetti *et al*., 2014; Neguembor *et al*., 2021; Fu *et al*., 2024).

Simulations suggest that the energy from transcription-generated supercoiling could facilitate TAD formation by displacing loaded cohesin complexes along chromatin, thereby promoting loop formation (Benedetti *et al*., 2014; Racko *et al*., 2018, 2019). Consistent with this prediction, in cells lacking the cohesin unloader, WAPL, cohesin accumulates at the 3’ end of genes, particularly at sites of convergent transcription (Busslinger *et al*., 2017). Further, in WAPL-deficient cells, excessive cohesin loading on chromatin produces microscopically visible nuclear structures called vermicelli (Tedeschi *et al*., 2013). These structures arise from unrestrained loop extrusion by cohesin (Haarhuis *et al*., 2017) and provide direct visual evidence that transcription-induced supercoiling can modulate loop formation (Neguembor *et al*., 2021). Although polymer simulations predict that elevated supercoiling within a TAD increases the frequency of contacts between genomic segments within the TAD, whether physiologically generated supercoiling accumulates within TADs *in vivo* – and measurably alters genomic contacts during cell differentiation – is yet to be demonstrated (Racko *et al*., 2019).

Here we report high-resolution mapping of DNA supercoiling during differentiation of pluripotent human embryonic stem cells to definitive endoderm, as a model of developmental commitment. This system combines experimental reproducibility with physiological relevance, enabling precise mapping of transcriptional and topological changes under the native regulatory control of the cell. By resolving supercoiling at multiple genomic scales, we demonstrate accumulation of negative supercoiling within highly transcribed genes and within enhancers, despite increased topoisomerase recruitment. Integration with high-resolution Hi-C maps to delineate 3D genome organisation, show that combined genic and enhancer RNAPII occupancy produces net negative supercoiling accumulation within A-compartment TADs (Lieberman-Aiden *et al*., 2009; Fortin and Hansen, 2015), which correlates with increased intra-TAD contact frequency. Together, these findings establish a genome-wide spatial and temporal link between transcriptional activity, supercoiling, and multi-scale genome organisation in human cells.

## Results

### A human stem cell model to study genome-wide changes in DNA supercoiling during differentiation

While previous research has predominantly emphasised the role of transcription factors and epigenetic regulators in directing cell fate transitions, the impact of supercoiling and its surveillance by topoisomerases, has been overlooked or proven difficult to study (Pommier *et al*., 2016; Björkegren and Baranello, 2018). The most used and best-characterised method for mapping DNA supercoiling is based on the intercalation of 4,5’,8-trimethylpsoralen (TMP) (Sinden *et al*., 1980, 1999). At limiting concentrations, TMP exhibits preferential binding to underwound as opposed to overwound DNA (Bermúdez *et al*., 2010) and can be crosslinked to the DNA with UV-A light (λ 365 nm) allowing for downstream enrichment of negatively supercoiled DNA. While TMP itself is cell permeable, commercially available derivatives carrying a biotin tag to allow enrichment prior to sequencing, are not. This necessitates the isolation and permeabilisation of nuclei prior to treatment (Krassovsky *et al*., 2021), which can affect the magnitude and distribution of supercoiling. Previous work mapping supercoiling using intercalation of cell-permeable derivatives of bTMP mainly relied on microarray for the analysis of genomic segments or specific promoters and were performed in simpler model organisms like yeast (Achar *et al*., 2020), *Drosophila* (Jupe *et al*., 1993) and *C. elegans* (Krassovsky *et al*., 2021), or was coupled to next-generation sequencing profiling cancer cell lines (Yao *et al*., 2025).

To explore how changes in transcription correlate with changes in genome supercoiling and 3D contacts, we exploited defined gene expression changes occurring during activation of developmental programs. We took advantage of a previously described and robust differentiation system in which H9 human embryonic stem cells are driven to the definitive endoderm germ layer (Teo *et al*., 2011; Yiangou *et al*., 2019; Eldridge *et al*., 2020) with high efficiency under defined conditions (Figure 1B). We performed RNA-seq with rRNA depletion at four time points between pluripotency (0 hours) and the end of the differentiation programme (72 hours), when >90% of the cells are definitive endoderm precursors (Figure S1A). Principal component and unsupervised clustering analysis of the top 500 most variable genes confirmed the reproducibility of the differentiation protocol and showed the distinct transcriptional signatures of pluripotent (0h), mesendoderm (24h) and endoderm (72h) cells (Figure 1C, D & S1B). As expected, key transcription factors involved in regulation of pluripotency maintenance and endoderm differentiation (Vallier *et al*., 2009; Teo *et al*., 2011), including POU5F1 (OCT4), SOX17, EOMES and GATA6 were among the top differentially expressed genes over the time course of the experiment (Figures 1E & S1C; Table S1).

To map changes in supercoiling across the genome, we synthesised a cell permeable psoralen moiety (biotinylated trimethyl psoralen, or bTMP), (Saffran *et al*., 1988) (Figure S2A, B & Methods) that has been previously used in conjunction with array hybridisation to map supercoiling in human chromosome 11 (Naughton *et al*., 2013) and in yeast (Achar *et al*., 2020). Briefly, human H9 cells were treated with bTMP (20 minutes, 37 °C) when undifferentiated (t = 0h) and when differentiated to definitive endoderm (t = 72h). Biotin TMP was crosslinked to DNA by exposing cells to 365 nm UV light and the excess molecule was washed out. bTMP-bound DNA was enriched on streptavidin beads before library preparation and next generation sequencing. The bTMP-seq signal is calculated by the genomic distribution of the log_2_ ratio of bTMP-enriched cellular DNA over the input (*i.e.* DNA that has been bTMP exposed but not subject to streptavidin pulldown). This signal is adjusted for sequence bias of bTMP incorporation by subtracting the log_2_ ratio of bTMP-enriched naked DNA to input (*i.e.* purified genomic DNA that has been exposed to bTMP but not enriched by streptavidin pulldown). A higher bTMP-seq signal represents greater negative supercoiling (Figure S2C).

To assess whether the concentration of bTMP affects the identification of supercoiling domains, we performed the experiment with 50 and 150 µg.ml^-1^ bTMP, with three biological replicates for each. The supercoiling signals for the combined data were strongly correlated (ρ >0.93) for both t = 0h and t = 72h samples (Figure S2D) showing that psoralen binding is independent of dose within the range employed.

We first inspected the supercoiling signal at individual genes exhibiting the greatest change in expression through differentiation, exemplified by OCT4, a pluripotency factor (Niwa *et al*., 2000) and SOX17, a defining transcription factor for definitive endoderm (Teo *et al*., 2011). RNA-seq confirmed downregulation of OCT4 and upregulation of SOX17, as expected (Figure 1E). Comparing the bTMP-seq signal through differentiation reveals a clear decrease in signal around OCT4 and an increase around SOX17 (Figure 1E), consistent respectively with dissipation and accumulation of negative supercoiling at these loci. Genome wide, an increase in negative supercoiling, as assessed by bTMP-seq in the bodies of long genes between the pluripotent (0h) and differentiated state (72h), correlates with an increase in gene expression (Figure 1F). This suggests that activation of genes during lineage commitment drives accumulation of underwound DNA at transcribed loci.

### Negative supercoiling in genes accumulates as a function of RNAPII occupancy despite increased topoisomerase recruitment

To explore the determinants of negative supercoiling accumulation in genes more detail we mapped genomic occupancy of transcriptionally engaged RNAPII using ChIP-seq with an antibody specific to the phosphorylated C-terminal domain of the Rpb1 subunit of RNAPII, and that of topoisomerases I, IIa and IIb (TOP1, TOP2α, TOP2β) using CUT&RUN. We first examined the correlation between bTMP signal enrichment (i.e. negative supercoiling) and transcriptional activity in ground-state pluripotent H9 ESCs. We segregated genes into four expression states based on RNA-seq signal. As expected, metagene plots show RNAPII occupancy is increased in the medium and highly expressed groups of genes (Figure 2A). Consistent with earlier studies (Kouzine *et al*., 2004, 2013; Naughton *et al*., 2013; Teves and Henikoff, 2014; Achar *et al*., 2020; Krassovsky *et al*., 2021), the bTMP signal (indicating negative supercoiling) is highest around the 5’ end of genes, tailing off to below baseline towards the 3’ end of the gene (Figure 2B). This general pattern is independent of differentiation state (Figure S3A) and bTMP dose (Figure S3B & C). Dividing the genes into groups by length clearly shows that the accumulation of negative supercoiling is confined to the 5’ segment of long genes, with the signal decreasing to baseline within approximately the first 5-10kb (Figure S3D). This is consistent with the twin supercoiling domain model (Liu and Wang, 1987) and the accumulation of positive supercoiling in the 3’ segment and TTS of long genes. The data also suggest that gene TSSs serve as barriers to the dissipation of negative supercoiling, but that negative supercoiling does spill into the surrounding chromatin at the TTS. While the bTMP assay only measures the degree of negative supercoiling, requiring positive supercoiling to be inferred, this is in line with recent observations in GM12878 cells using azide-TMP (Yao *et al*., 2025) and studies mapping positive supercoiling by means of GapR binding (Guo *et al*., 2021; Longo *et al*., 2024).

**Figure 2.**
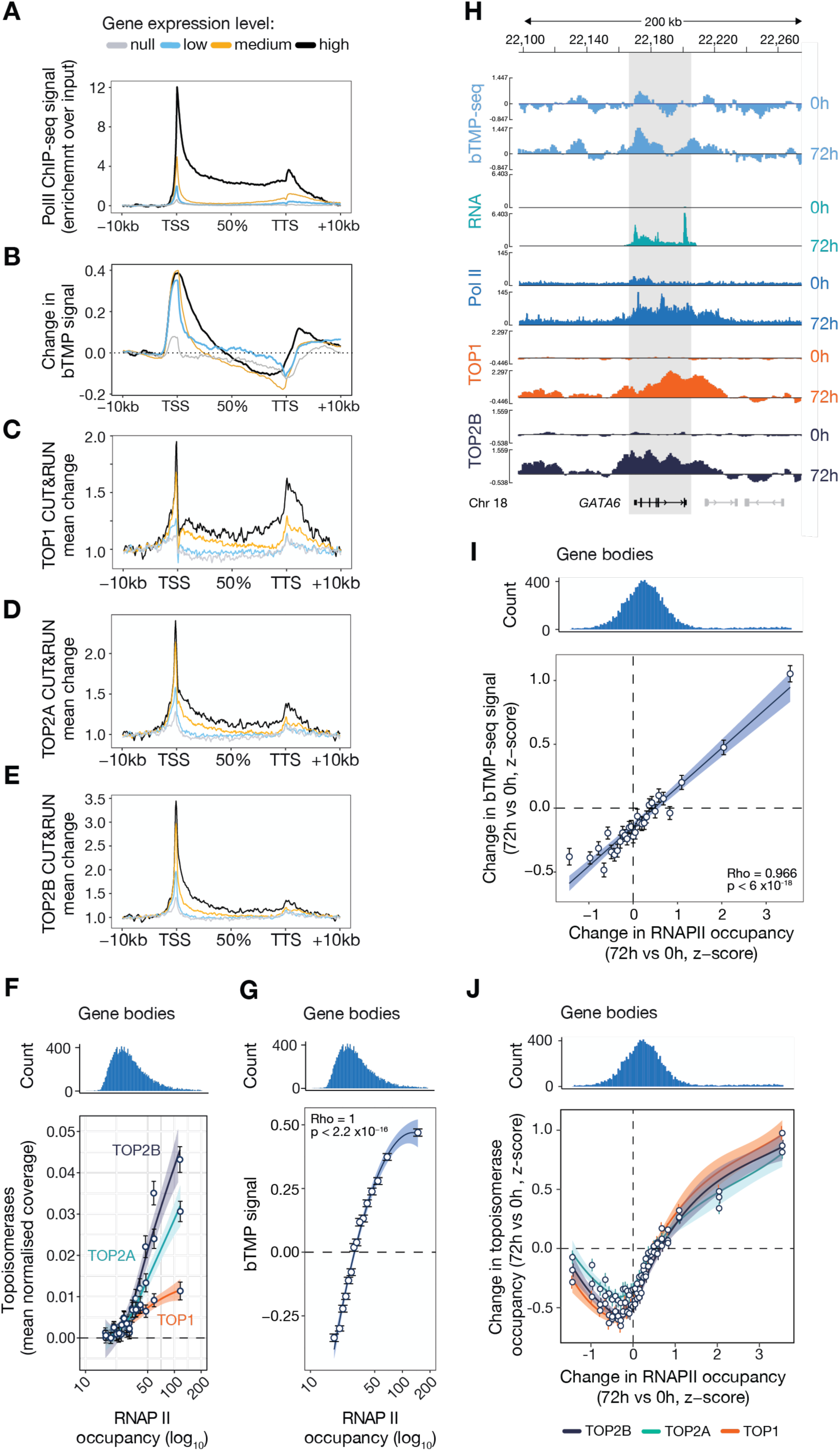
Topoisomerases are recruited as a function of transcription but do not fully control negative supercoiling. A. Metagene analysis of RNAPII occupancy in genes >1kb long in undifferentiated H9 cells binned into four groups by gene expression determined by RNA-seq. For panels A – E, the expression groups are: no expression – light grey; low (quantile < 0.5), blue; medium (quantile = 0.5 – 0.9), orange; and high (quantile > 0.9), dark grey. Coverage matrices for each category were computed at 50 bp-window resolution except the bTMP signal, which was computed at 500 bp windows. Mean changes (enrichment over background) were computed in 400 bins (100 in each flank, 200 in gene) and normalised using background values from domains adjacent to genes (−10 to –5 kb). TSS = transcription start site; TTS = transcription termination site. B. Metagene of normalised bTMP incorporation in undifferentiated H9 cells by expression group. C. Metagene of topoisomerase 1 occupancy in undifferentiated H9 cells by expression group. D. Metagene of topoisomerase 2a occupancy in undifferentiated H9 cells by expression group. E. Metagene of topoisomerase 2b occupancy in undifferentiated H9 cells by expression group. F. Correlation between RNAPII signal and mean TOP1 (orange), TOP2α (green) and TOP2β (dark purple) enrichment at gene bodies from coding genes that are longer than 3 kb. Gene bodies exclude 1 kb downstream of the TSS and 1 kb upstream of the TTS. Background regions were defined as –10 to –5 kb upstream of a gene. Data points and error bars represent the median CUT&RUN enrichment ± SEM. Spearman correlation and p values: TOP1 ρ = 0.899, p = 2.65 x10^-7^; TOP2α ρ = 0.733, p = 7.64 x10^-6^; TOP2β ρ = 0.716, p = 1.52 x10^-5^. G. Correlation between median supercoiling score (relative to background) and RNAPII signal in gene bodies from coding genes longer than 3 kb. Gene bodies exclude 1 kb downstream of the TSS and 1 kb upstream of the TTS. Background regions were defined as –10 to –5 kb upstream of a gene. Data points and error bars represent the mean supercoiling score ± SEM. RNAPII occupancy normalised to 1 kb was defined as the sum (score over gene body / mean length of the gene body) x 1000. H. Representative sequencing tracks of bTMP, RNAPII, TOP1 and TOP2β signal at the GATA6 locus (grey zone) in pluripotent (0h) and definitive endoderm (72h) cells. Top scale: genomic coordinates on chromosome 18. I. Pearson correlation between changes in supercoiling (ΔbTMP-seq signal, z-score) and RNAPII occupancy within gene bodies at 0 *vs*. 72 hours. J. Correlation between changes in topoisomerase recruitment (z-score) and RNAPII occupancy (z-score) within gene bodies between 0 and 72 hours. All curves fitted with LOESS.

Both type I (TOP1) and type II (TOP2α & TOP2β) topoisomerases are enriched at transcriptionally active genes (Figure 2C – E) and accumulate as a function of RNAPII occupancy (Figure 2F). However, dissecting the distinct contributions of each enzyme to supercoiling control is challenging. While previous studies have suggested a major role of TOP1 and TOP2B, but not TOP2A, in the relaxation of the transcription-dependent supercoiled DNA (Yao *et al*., 2025), our immunoblot and RT-qPCR data show that the TOP2A mRNA and protein levels are higher than TOP2B in pluripotent H9 cells (t = 0h) (Figure S3E & F). TOP2A is then downregulated during endoderm lineage commitment as TOP2B expression increases. However, while topoisomerase recruitment increases as a function of transcriptional activity, so does net negative supercoiling; despite greater overall topoisomerase recruitment, negative supercoils are generated faster than they can diffuse or be enzymatically relaxed (Figure 2G). Note that at high levels of RNAPII occupancy, the bTMP-seq signal appears to saturate. This may reflect a supercoiling maximum beyond which further negative superhelicity cannot be accommodated, although we cannot exclude that it may also reflect a limit to the dynamic range of the assay.

To examine the dynamics of supercoiling during endoderm lineage commitment we next compared the bTMP signal in cells at pluripotency (0h) and definitive endoderm (72h), alongside RNAPII and topoisomerase occupancy. Taking as an example the GATA6 locus, a transcription factor upregulated on endoderm differentiation (Fisher *et al*., 2017), productive RNAPII accumulation across the gene body by 72h is accompanied by an increase in the bTMP signal and recruitment of topoisomerases (Figure 2H). Across the genome, changes in RNAPII levels in gene bodies between 0h and 72h correlated closely with changes in supercoiling as genes are up and down regulated (Figure 2I), broadly in line with the correlation between supercoiling and changes in mRNA levels (Figure 1F). The greatest positive changes in RNAPII between the two time points lead to the greatest increases in topoisomerase recruitment (Figure 2J), consistent with demand-driven recruitment of topoisomerases to resolve the increased torsional stress that accompanies increased transcription (Figure 2I). In contrast, at genes in which RNAPII occupancy is downregulated, decreases in topoisomerase recruitment were less clearly proportional to RNAPII loss. This asymmetry suggests that topoisomerase recruitment is dynamically upregulated in response to increased torsional flux at genes activated during differentiation. In contrast, during gene repression, changes in net supercoiling, although remaining monotonic with RNAPII occupancy (Figure 2I), are primarily driven by reduced torsion generation and/or altered topoisomerase.

We next asked whether negative supercoiling accumulates at sites of non-genic transcription, specifically at functional enhancers, which are transcribed by RNAPII into enhancer RNAs (eRNAs) (Kim *et al*., 2010; Koch *et al*., 2011). We mapped active enhancers in pluripotent H9 cells according to previously defined criteria (Lyu *et al*., 2018) as loci enriched in H3K4me1 that overlap with open chromatin (ATAC-seq peaks), but are not enriched for H3K4me3 or occur within 1kb of TSSs. ‘Meta-enhancer’ plots in which the identified enhancers were binned by RNAPII occupancy demonstrate a higher bTMP-seq signal in those enhancers with the highest levels of RNAPII occupancy (Figure 3A & E), accompanied by enhanced topoisomerase recruitment, particularly of type II topoisomerases (Figure 3B-D, F – H). Thus, enhancers exhibit a similar relationship between transcription, supercoiling and topoisomerase recruitment as observed in genes, reinforcing the similarity and overlap between enhancers and promoters (Andersson and Sandelin, 2020).

**Figure 3.**
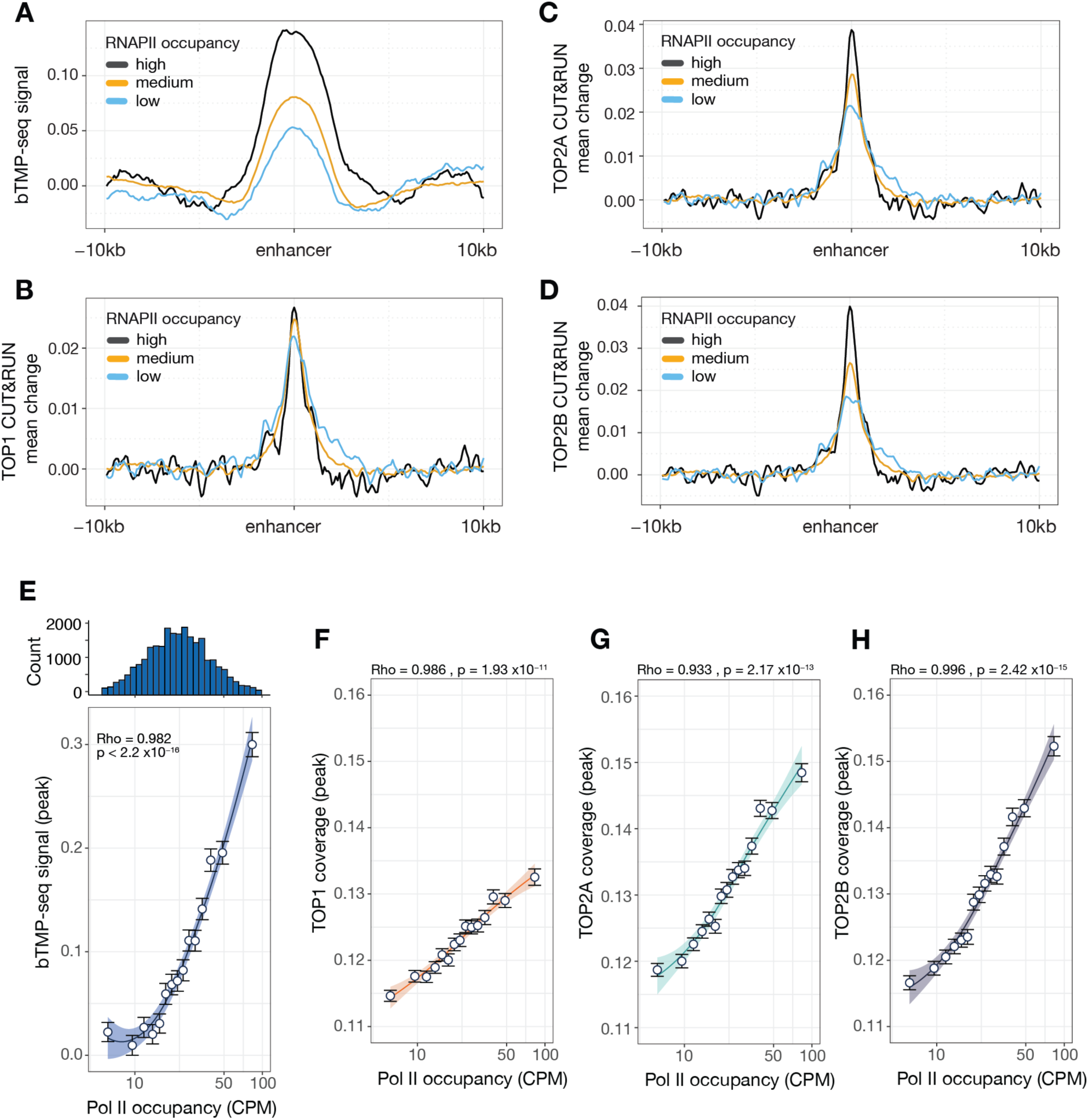
Transcription induced supercoiling at enhancers in undifferentiated H9 cells. A. Supercoiling distribution in enhancers as a function of enhancer RNAPII occupancy. Enhancer RNAPII was binned into three groups, light grey; low (quantile < 0.1), blue; medium (quantile = 0.1 – 0.9), orange; and high (quantile > 0.9), dark grey. Supercoiling coverage matrices were computed at 50 bp-window resolution for topoisomerases and 500 bp for bTMP-seq signal. Mean change in supercoiling (enrichment over background) was computed in 200 bins and normalised to background using values from domains adjacent to enhancers: –10 to –5 kb and +10 to +5 kb. B – D). Topoisomerase enrichment mapped by CUT&RUN at stitched enhancers as a function of eRNA levels in H9 cells. B. TOP1; C. TOP2α; D. TOP2β. E. Correlation between normalised bTMP-seq supercoiling signal and RNAPII occupancy. The distribution of observations is shown above the graph. Each point represents the mean peak SS signal within enhancers grouped into 15 bins according to their Pol II occupancy (CPM). Error: SEM of SS peak values within each Pol II bin. Curve fit: LOESS; correlation: Spearman’s ρ. F – H. Correlation between normalised topoisomerase coverage (F. TOP1; G. TOP2A; H. TOP2B) and RNAPII occupancy. Data points represent the mean peak topoisomerase signal within enhancers grouped into 15 bins according to their Pol II occupancy (CPM). Error: SEM of SS peak values within each Pol II bin. Curve fits: LOESS; correlation: Spearman’s ρ.

### Negative supercoiling correlates with increased contacts within gene bodies

While transcription-induced supercoiling generates plectonemes, which increase DNA contacts in single-molecule supercoiling assays (Janissen *et al*., 2024), the extent to which this occurs in chromatinised domains in cells, and whether supercoiling can drive intragenic contacts remains less clear. To address this question, we generated Hi-C maps of H9 cells in their pluripotent state (0 hours) and after 72 hours of endoderm differentiation. Hi-C interaction maps centred on TSS and termination sites (TTS) show a clear increase in intragenic contacts as a function of expression (Figure 4A) or RNAPII binding (Figure 4B) in undifferentiated cells. For highly expressed genes, we observed interactions between the TSS and TTS that were dynamically gained or lost as genes are activated or repressed during differentiation, respectively (compare upper and lower panels of Figure 4C & 4D). These observations indicate that RNAPII engagement promotes local chromatin interactions, potentially through transcription-induced negative supercoiling, though recruitment of multiple RNAPII complexes and Mediator– or cohesin-dependent bridging (Kagey *et al*., 2010; Mattingly *et al*., 2022; Ramasamy *et al*., 2023) may also underlie this effect. Consistent with this idea, RNAPII genome-wide enrichment strongly correlated with both negative supercoiling and insulation score at 0 and 72h (Figure 4E). The insulation score, derived from Hi-C data, reflects the number of contacts made by a given genomic segment (Crane *et al*., 2015). Together, these findings are consistent with previous observations showing that elongating RNAPII promotes gene loop formation (O’Sullivan *et al*., 2004; Tan-Wong *et al*., 2012; Rowley *et al*., 2019) and suggest that transcription-induced negative supercoiling may provide a physical mechanism linking RNAPII engagement to enhanced intragenic chromatin interactions.

**Figure 4.**
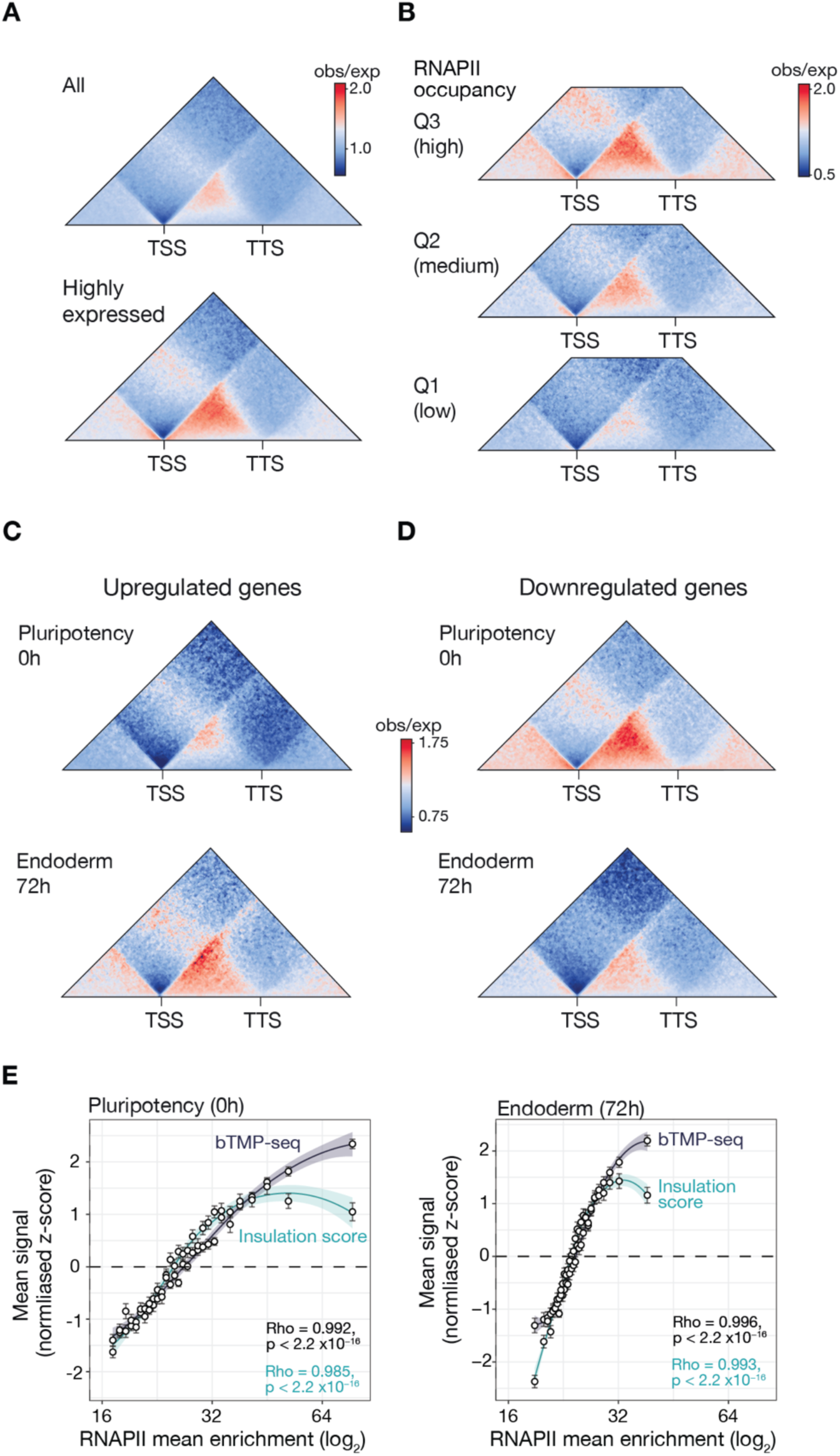
Transcription induced supercoiling correlates with increased intragenic contacts. A. Aggregate normalised contact matrices between the TSS and TTS of all coding genes (upper panel) and highly expressed coding genes only (lower panel). B. Aggregate normalised contact matrices between the TSS and TTS of coding genes binned by RNAPII occupancy. C. Analysis of intra-gene normalised interactions between the TSS and the TTS of genes upregulated (p ≤ 0.05) between 0h and 72h of endoderm differentiation. D. Analysis of intra-gene normalised interactions between the TSS and the TTS of genes downregulated (p ≤ 0.05) between 0h and 72h of endoderm differentiation. E. Correlation of insulation score (green) and bTMP-seq signal (dark purple) with RNAPII mean enrichment at 0h and 72h. Spearman’s ρ.

### TADs within compartment A exhibit greater negative supercoiling than those in compartment B which correlates with increased intra-TAD contacts

We next asked whether the principles observed at the level of individual genes extend to higher order genome organisation, specifically within topologically associating domains (TADs). TADs are considered fundamental units of gene regulation (Bonev and Cavalli, 2016; Zheng and Xie, 2019), and determining whether they are supercoiled is an open question that could inform how enhancer-promoter communication is established (Racko *et al*., 2019).

Using a window of 25 kb, we identified 5,603 TADs with an average width of ∼500 kb, evenly distributed between the A and B compartments. A representative genomic region illustrating RNA levels, supercoiling and genome contacts (insulation score and TADs) in the context of computationally defined A and B compartments is shown in Figure 5A. Analysis of average negative supercoiling in undifferentiated cells (t = 0 h) revealed that TADs in compartment A exhibit substantially greater negative supercoiling than those in compartment B (Figure 5B & C). The borders of compartment A TADs correspond to negative supercoiling maxima, a pattern absent in TADs of compartment B (Figure 5B). The same patterns were observed in differentiated cells (Figure S4A & B).

**Figure 5.**
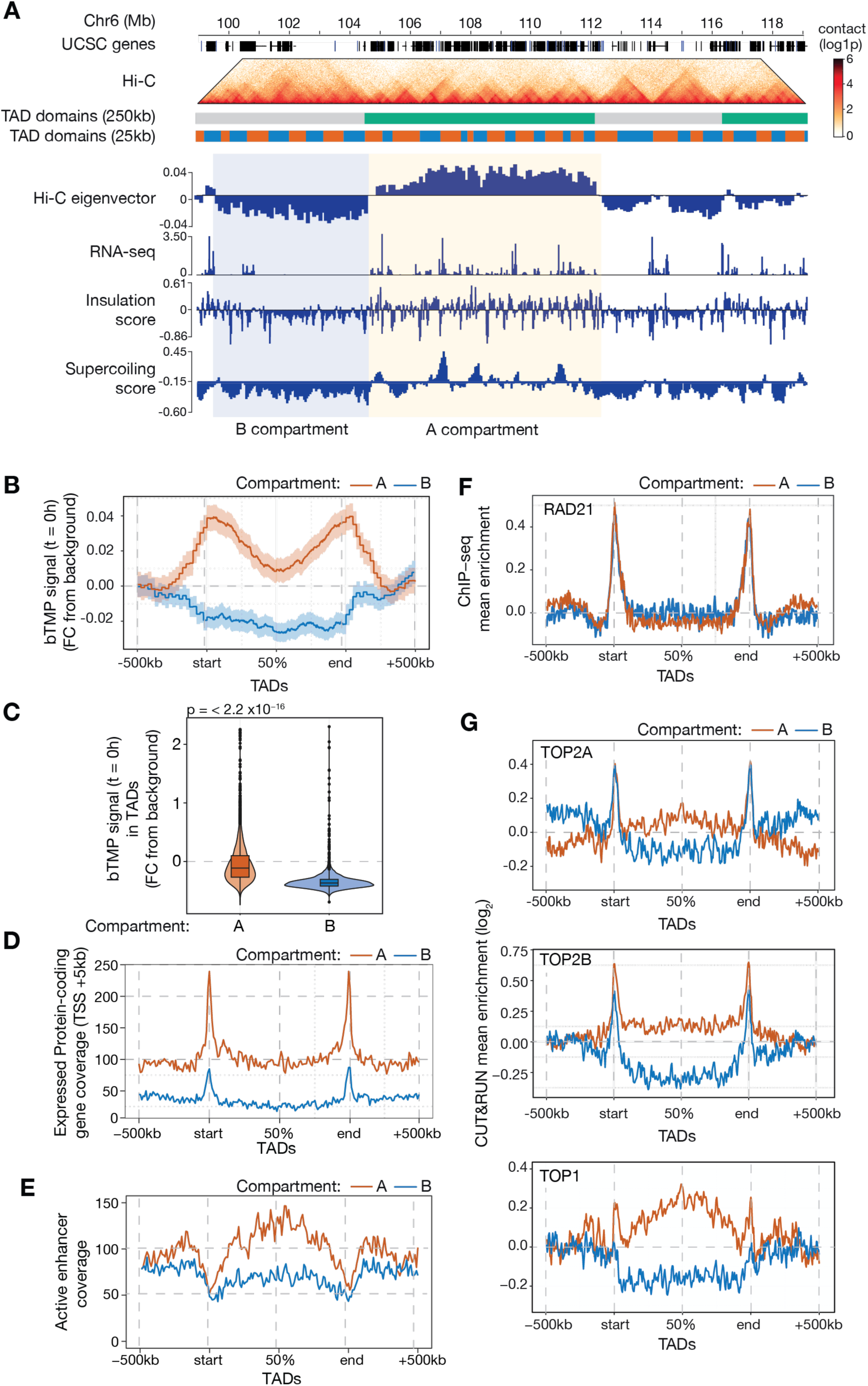
Supercoiling in TADs. A. Example genomic tracks displaying the Hi-C contact matrix with TADs called with 250 kb and 25 kb windows; the Hi-C eigenvector, which defines chromosomal compartments (example A compartment is shaded in yellow; example B compartment is shaded in blue), insulation score, RNA-seq and bTMP-seq normalised signals across a 20 Mb interval of chromosome 6 of human H9 cells. UCSC genes are displayed in black boxes and genomic coordinates are shown at the top. B. bTMP-seq signal variation normalised to background within TADs at t = 0h binned by compartment (A = red; B = blue). TAD boundaries are denoted as ‘start’ and ‘end’ and the middle meta-distance is denoted as 50%. C. Quantitation of bTMP-seq signal in TADs of the A (red) and B (blue) compartments at t = 72h. P value calculated with a two-sample Kolmogorov-Smirnov test. D. Distribution of expressed (RNAPII Q2 & Q3 occupancy) protein coding genes (TSS + 5kb) in TADs binned by compartment (A = red; B = blue). E. Active enhancer distribution in TADs of the A compartment (red) and B compartment (blue). F. RAD21 enrichment in TADs of the A compartment (red) and B compartment (blue). G. Distribution of TOP2α, TOP2β and TOP1 in meta-TADs of compartment A (red) and B (blue).

To identify potential sources of negative supercoiling in compartment A TADs, we analysed the distribution of genes and enhancers. Since negative supercoiling is strongest within the first 5 kb downstream of TSSs (Figure S3D), we focused on this region for gene-associated signal. Protein-coding genes are enriched in compartment A relative to compartment B (16,717 vs 7,287) and preferentially localise near TAD boundaries (Figure 5D), consistent with the elevated negative supercoiling observed at A-compartment TAD borders. We next assessed enhancer contributions by classifying previously identified enhancers (Figure 3) as active, primed, or poised based on histone modifications (reviewed in Spicuglia and Vanhille, 2012). As expected, active enhancers engage in more long-range chromatin contacts than primed or poised enhancers (Figure S5) and are preferentially enriched in compartment A (Figure 5E). Within A-compartment TADs, active enhancers are concentrated toward the TAD centre, whereas they are not enriched in B-compartment TADs (Figure 5E). Although enhancers generate less negative supercoiling than genes (compare the magnitude of the bTMP signal between enhancers in Figure 3A and genes in Figure 2B), their enrichment in A-compartment TAD bodies likely contributes to the elevated supercoiling in the centre of A-compartment TADs relative to those in the B-compartment. As expected, the cohesin subunit RAD21 accumulates at TAD boundaries in both compartments (Figure 5F and Dixon *et al*., 2012 #107454; Rao *et al*., 2014 #173635). Type II topoisomerases are also enriched at TAD boundaries irrespective of compartment, and additionally throughout the body of A-compartment TADs (Figure 5G), in agreement with previous studies (Uusküla-Reimand *et al*., 2016; Canela *et al*., 2019). In contrast, TOP1 is enriched at the boundary and interiors of compartment A TADs (Figure 5G).

Together, the spatial arrangement of genes and enhancers within A-compartment TADs establishes a supercoiling gradient, with torsional stress increasing from the TAD centre toward its boundaries. This organisation provides a plausible physical basis for directional loop extrusion and suggests that transcription-driven supercoiling acts in concert with CTCF and cohesin to reinforce the asymmetric architecture of TAD loops.

*In silico* modelling has suggested that the accumulation of negative supercoiling within transcriptionally active TADs promotes enhancer – promoter communication and may facilitate promoter opening, analogous to the action of DNA gyrase in bacteria (Racko *et al*., 2018, 2019). In agreement with this model, and extending our gene-level observations, the higher levels of negative supercoiling displayed by TADs in compartment A (Figure 5B) is accompanied by an increased number of normalised chromatin contacts and more strongly insulated boundaries relative to compartment B TADs (Figure 6A – C). Across both pluripotent cells (0 h) and endoderm precursors (72 h), genome wide negative supercoiling shows a strong correlation with insulation score (Figure 6D & E), confirming that elevated torsional stress is associated with enhanced contact frequency.

**Figure 6.**
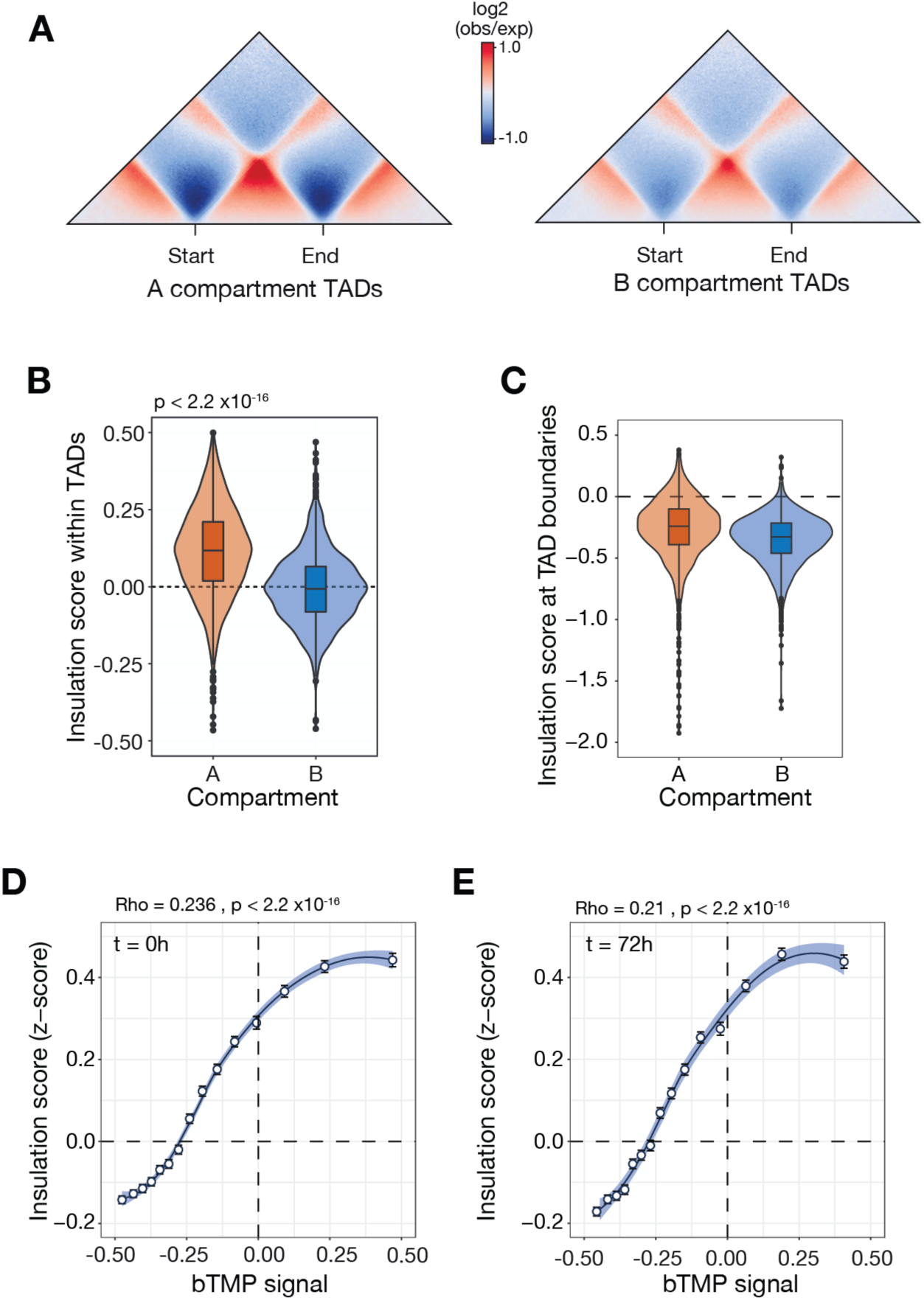
Negative supercoiling in TADs correlates with increased intra-TAD. A. Normalised contact matrices for TADs in compartment A and compartment B. B. Mean insulation score within TADs in compartment A (red) and compartment B (blue). P value calculated with a two-sample Kolmogorov-Smirnov test. C. Mean insulation score at the boundaries of TADs in compartment A (red) and compartment B (blue). P value calculated with a two-sample Kolmogorov-Smirnov test. D & E. Correlation between supercoiling signal and insulation score (z-score) in TADs at 0h (D) and 72h (E). Data is divided into 15 bins with the mean and standard error of the mean indicated for each. Correlation test: Spearman ρ.

### Higher order organisation of supercoiling and TADs in the human genome

Finally, we asked whether supercoiling varies across higher levels of genome organisation. We identified TADs at a coarser Hi-C resolution of 250 kb, defining 712 larger domains (’SuperTADs’) with an average size of ∼4.5 Mb that collectively encompass the 5,531 TADs previously identified at 25 kb resolution. To examine how individual TADs are arranged within these larger structures, we generated aggregate profiles in which each TAD was positioned relative to its host SuperTAD (Figure 7A). As expected, CTCF and the cohesin subunit RAD21 are enriched at individual TAD boundaries, consistent with their roles in loop anchoring and domain insulation. Strikingly, however, TADs located at SuperTAD boundaries exhibit a higher density of protein-coding genes, increased RNAPII occupancy and mRNA levels, and correspondingly elevated negative supercoiling compared with internal TADs (Figure 7A). These boundary TADs are also depleted for contacts with LaminB1 mapped by DamID (Pickersgill *et al*., 2006; Meuleman *et al*., 2013), indicating that they are positioned further from the nuclear lamina and deeper within the nuclear interior.

**Figure 7.**
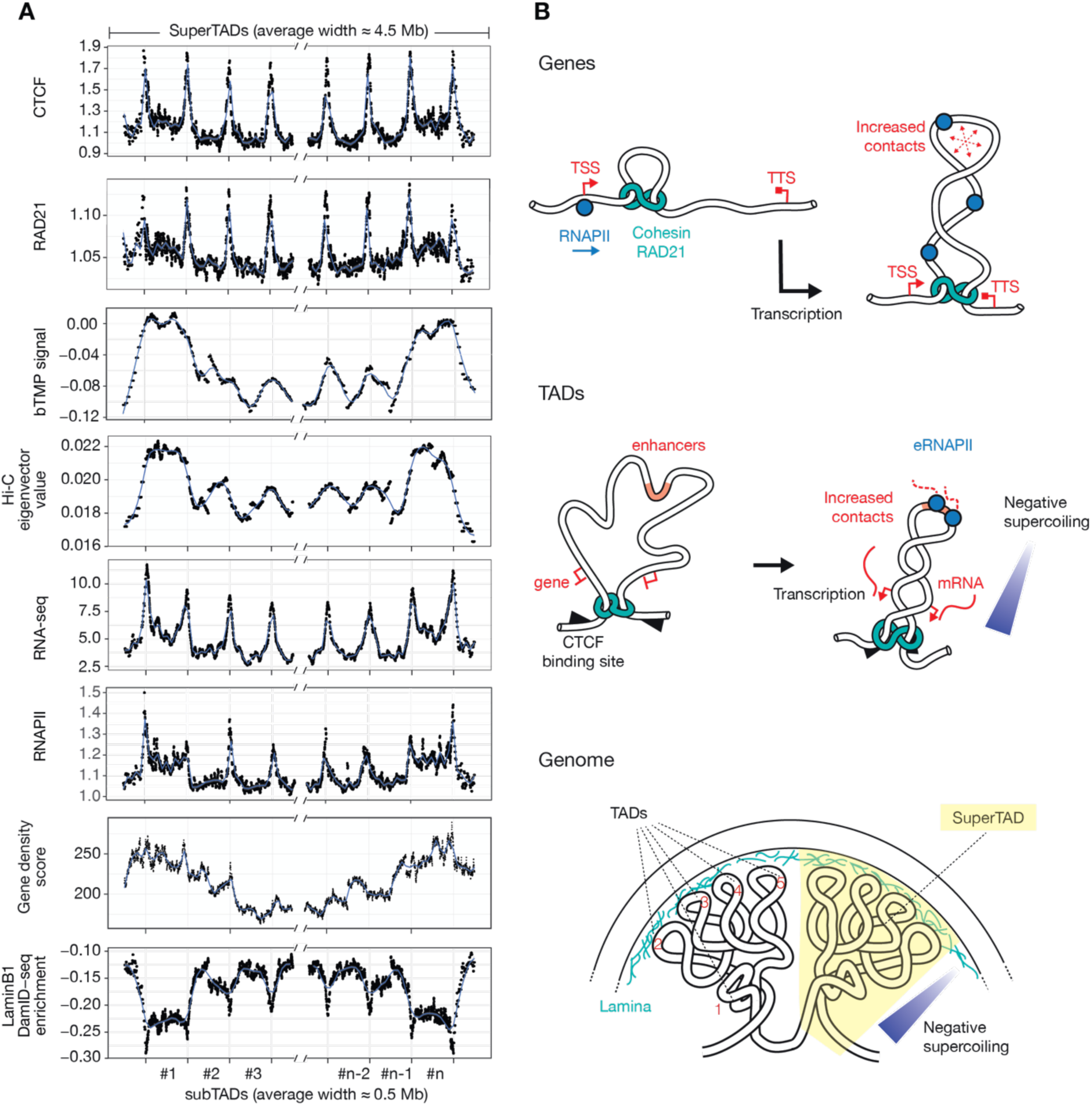
Negative supercoiling defines a larger scale organisation of TADs. A. TADs called with a 25kb window aligned within their corresponding ‘SuperTAD’ called with a 250kb window. Although there are, on average ∼10 TADs per SuperTAD domain, the figure shows just three from each end of the SuperTAD, labelled #1 – 3 and #n – #n-2. Features mapped: CTCF and RAD21 ChIP-seq, supercoiling, the Hi-C eigenvector value, mRNA measured by RNA-seq, RNAPII occupancy and gene density. The LaminB1 DamID-seq is from the SHEF2 human embryonic stem cell (Meuleman *et al*., 2013), GEO accession number GSM557443. B. Supercoiling and genome contacts at multiple scales. Top panel: genes. At the scale of genes, transcription-induced supercoiling leads to increased contacts within gene bodies and between the TSS and TTS, which may reflect the formation of plectonemic structures within the gene, driven by negative supercoiling. Middle panel: TADs. Transcriptionally active, compartment A, TADs exhibit more negative supercoiling than those with low levels of transcription. This correlates with higher levels of contacts, which may facilitate, for example, enhancer-promoter interactions. The accumulation of supercoiling within a TAD may be driven by both enhancer transcription and gene transcription, enhancers tending to be found at the core of the TAD, while genes are more likely to be found at the extremities orientated towards the base of the TAD. Consistent with the proposal of Racko et al., enhancer transcription scales with negative supercoiling within TADs and may facilitate promoter melting by contributing to a net negatively supercoiled environment in the TAD. Lower panel: genome. The unexpected behaviour of smaller TADs at the extremities of large-scale ‘SuperTADs’ (TAD number 1 in the figure) suggests that they behave differently from their neighbours (TADs 2-5). This suggests the possible organisation of genome loops depicted with SuperTADs comprising a discrete group of TADs, the outer of which are more centrally placed and more negatively supercoiled than TADs at the core of the SuperTAD, which are more likely to interact with the nuclear lamina.

Together, these observations support a model in which groups of transcriptionally active TADs positioned away from the nuclear envelope flank lamina-associated chromatin to form a higher-order, multi-TAD structure that we term a SuperTAD (Figure 7B).

## Discussion

Here we have used stem cell differentiation as a tool to drive physiologically relevant changes in gene expression to explore the interplay between transcription-driven DNA supercoiling and 3D genome contacts. Our observations support the principle that, across multiple genomic scales, increased steady-state negative supercoiling is generated by transcription and, in turn, is associated with an increased frequency of contacts within a domain.

At the level of individual genes, our data indicate that the TSS acts as at least a partial barrier to supercoil diffusion, as negative supercoiling does not spread upstream of the TSS (Figure 2B and S3A–D). This may reflect the presence of cohesin at active promoters (Busslinger *et al*., 2017; Valton *et al*., 2022), enabling the accumulation of negative supercoiling within the gene body in proportion to RNAPII occupancy. This behaviour is consistent with the twin-domain model (Liu and Wang, 1987), in which transcription generates positive supercoiling ahead of elongating RNAPII and negative supercoiling behind it. Preferential resolution of positive supercoiling, proposed to be mediated by TOP1 (Baranello *et al*., 2016) and TOP2 (Fernández *et al*., 2014), would not only mitigate a topological barrier to transcription elongation (Ma *et al*., 2013) but also effectively confine negative supercoiling to the gene body downstream of the TSS. In long genes (Figure S3D), the gradient of supercoiling between the 5′ and 3′ ends is more pronounced, with the previously reported dip toward the transcription termination site (TTS) clearly visible (Yao *et al*., 2025), consistent with the accumulation of positive supercoiling. Together, these findings suggest that although topoisomerases are recruited to active genes, continuous RNAPII elongation produces negative supercoiling faster than it can be resolved, particularly in short, highly transcribed genes, which therefore act as potent local sources of torsional stress on the surrounding chromatin.

A simple explanation for the link between negative supercoiling and increased genomic contacts within genes, consistent with extensive theoretical and experimental work on naked DNA (Fogg *et al*., 2021; Pyne *et al*., 2021; Janissen *et al*., 2024), is that torsional energy stored in negatively supercoiled DNA between the TSS and TTS is converted into folding of the chromatin fibre into plectonemic supercoils, analogous to writhe in naked DNA. This folding would generate contacts within gene bodies and between the TSS and TTS (Figure 4). Such gene loops have been proposed to facilitate RNAPII recycling, sustain high transcriptional output and enable rapid regulation of gene activity (Rowley *et al*., 2017, 2019).

At the level of TADs, we observe a similar relationship: accumulation of negative supercoiling within A-compartment TADs correlates with increased intra-TAD contacts. These observations are broadly consistent with computational modelling predicting that active (compartment A) TADs are supercoiled (Benedetti *et al*., 2014; Racko *et al*., 2018), and that such supercoiling promotes intradomain contacts (Racko *et al*., 2018) and facilitates promoter opening (Racko *et al*., 2019), in a manner analogous to the role proposed for DNA gyrase in regulating certain E. coli promoters (Smith *et al*., 1978).

The presence of a supercoiling gradient across active TADs raises the question of how torsional stress can act over such genomic distances. Previous work suggests that diffusion of negative supercoiling away from transcription units is limited and does not extend far beyond the gene itself (Kouzine *et al*., 2013). Consistent with this, we observe that negative supercoiling drops sharply at the 5′ end of genes and extends no more than ∼5 kb beyond the 3′ end, potentially up to ∼10 kb in short, highly transcribed genes (Figures 2B and S3D). However, it is important to note that our analyses rely on population-averaged measurements; supercoiling states are likely to be highly dynamic and heterogeneous at individual loci, and averaging may therefore underestimate the true extent or variability of torsional stress in single cells. Regardless of the precise diffusion distance, the cumulative activity of many transcription units, each acting as a local torsional ‘motor’, can generate a net accumulation of negative supercoiling within active TADs.

This raises the possibility that supercoiling gradients contribute to, or modulate, cohesin-mediated loop extrusion. Recent modelling and single-molecule experiments suggest that DNA torsional stress can influence cohesin and SMC complex translocation, loop size, and stability (Benedetti *et al*., 2017; Racko *et al*., 2018; Neguembor *et al*., 2021; Davidson *et al*., 2023; Jeppsson *et al*., 2024). Within active TADs, negative supercoiling may promote local chromatin folding and bias cohesin movement, potentially preventing reversal and providing a ratchet-like mechanism that reinforces directional loop extrusion. In this way, supercoiling and loop extrusion could act in concert to stabilise TAD architecture and facilitate enhancer–promoter communication. Increased contacts within TADs may further reinforce transcription by promoting interactions between genes and their regulatory elements, establishing a feedforward mechanism for robust gene activation (Deng *et al*., 2012; Beagrie *et al*., 2017; Lyu *et al*., 2018).

Finally, our data reveal a previously unappreciated level of higher-order organisation within TADs (Figure 7B). By grouping TADs identified at a resolution of 25 kb into larger ‘SuperTADs’ detected at 250 kb resolution, we find that TADs located at the edges of these domains exhibit distinct properties: they are more gene-dense, more negatively supercoiled, and likely positioned deeper within the nuclear interior. This hierarchical organisation may shape the distribution of torsional stress across the genome and thereby influence local chromatin function. Taken together, our observations identify DNA supercoiling as a dynamic, transcription-coupled determinant of genome organisation, linking gene activity to chromatin folding and 3D contacts across multiple scales.

## Materials & Methods

### Human embryonic stem cell culture

The human embryonic stem cell line WA09/H9 (XX karyotype) was obtained from the Lancaster lab (MRC LMB) and used under WiCell Research Institute agreement 20-W0364. Cells were cultured in Essential 8 Flex medium (Gibco) on Vitronectin-XF (STEMCELL Technologies) coated 6-well plates at 37°C, 5% CO_2_. Cells were passaged every 3-4 days and diluted as desired based on cell density. For passaging, cells were washed and incubated for 5 minutes at room temperature in 0.5 mM UltraPure EDTA (Thermo Fisher Scientific), and gently detached from the plates by pipetting colonies while keeping them as small aggregates. Regular testing confirmed absence of mycoplasma contamination.

### Endoderm differentiation

Differentiation of H9 hESC to definitive endoderm precursors was performed as previously described (Teo *et al*., 2011). Differentiation was initiated by replacing Essential 8 Flex medium with CDM-PVA supplemented with 100 ng/mL Activin A (Hyvönen group, Biochemistry Department, University of Cambridge), 80 ng/mL FGF2 (R&D), 10 ng/mL BMP4 (R&D), 10 μM PI3K inhibitor LY294002 (Promega), and 3 μM GSK3 inhibitor CHIR99021 (Tocris). The next day, medium was changed to CDM-PVA supplemented with Activin A, FGF2, BMP4 and LY294002 as above. CDM-PVA comprised 50% (v/v) Ham’s F-12 (Gibco) and 50% (v/v) IMDM (Gibco) supplemented with 1 g/L polyvinyl alcohol (PVA, Sigma), 1X Chemically Defined Lipid Concentrate (Gibco), 0.5 mM Thioglycerol (Sigma), 15 μg/mL Transferrin (Roche) and 7 μg/mL Insulin (Roche). On day three, medium was replaced with RPMI+supplemented with 100 ng/mL Activin A and 80 ng/mL FGF2. RPMI+ was made up of RPMI with GlutaMax (Gibco), 1X MEM Non-Essential Amino Acids Solution (Gibco) and 1X B27 supplement serum-free, with insulin (Gibco). Differentiation efficiency of each experiment was monitored using intracellular and cell surface antigen detection by flow cytometry, employing antibodies targeting SOX17, EOMES, and CXCR4 (CD184). Antibodies used in the study are listed in Table S2. Flow cytometry analyses were performed using a BD Fortessa Cell Analyser (BD Bioscience).

### RNA extraction and sequencing (RNA-seq)

To isolate RNA, medium was aspirated from plates, adherent cells were washed twice in DPBS and lysed in 700 μL RLT lysis buffer containing β-mercaptoethanol (Sigma-Aldrich). Lysates were transferred to sterile tubes and stored at –80°C. RNA was extracted from all samples simultaneously by column purification using the Rneasy Kit (QIAGEN) according to the manufacturer’s protocol and quantified using a Nanodrop 1000 (Thermo Fisher Scientific). Libraries were constructed employing the NEBNext Ultra II RNA library preparation kit and NEBNext rRNA depletion kit (NEB). cDNA libraries were generated from 750 ng of RNA, amplified through 8 PCR cycles, and indexed with NEBNext Multiplex Oligos for Illumina (NEB). Quality control checks, including testing for primer and adapter contamination, were performed using the Agilent 2100 Bioanalyzer High Sensitivity DNA chip. Subsequently, libraries were quantified using a KAPA Library Quantification Kit (Roche), pooled, and subjected to high-throughput sequencing. Sequencing was performed in a 1×50 bp single read format using an Illumina HiSeq 4000 flow cell at the CRUK Genomics Core Facility.

### Chromatin immunoprecipitation sequencing (ChIP-seq)

Following removal of spent medium, cells were washed in DPBS, treated with pre-warmed StemPro Accutase (Gibco), and incubated at 37°C until primarily single cells were observed. Approximately 15-20 million cells were diluted in growth media and fresh formaldehyde was added to a final concentration of 1%. Cross-linking reaction was performed for 10 minutes at room temperature and stopped by adding glycine solution (0.25 M final concentration) for 5 minutes. Cells were then pelleted, washed in ice-cold PBS, and snap-frozen in liquid nitrogen (−135°C), for storage at –80°C. To obtain nuclei, cell pellets were thawed on ice and lysed by resuspending in cold lysis buffer (10 mM Tris-HCl pH 7.5, 10 mM NaCl, 3 mM MgCl2, 0.5% NP40), followed by incubation on ice for 5 minutes. Cells were spun down at 2000 g for 5 minutes, and steps were repeated twice for a total of three lysis. Crude nuclear extracts were immediately resuspended in sonication buffer (1% SDS, 10 mM EDTA, 50 mM Tris-HCl pH 8.0), aliquoted in Diagenode TPX tubes (Diagenode), and equilibrated on ice for 20-30 minutes with occasional vortexing. Chromatin was sonicated to an average size of 300-500 bp using a Bioruptor device with an integrated cooling system (Diagenode) for 30 cycles (30 seconds ON/30 seconds OFF; High power setting; 4°C). Chromatin Dilution Buffer (1.2% (v/v) Triton X-100, 22 mM Tris-HCl pH 7.5, 5 mM EDTA pH 8, 182 mM NaCl, 0.15% (w/v) Sodium deoxycholate) was added to sonicated samples and each tube was spun at 13,600 g for 30 minutes at 4°C to pellet any insoluble material. An aliquot from the supernatant containing soluble chromatin was taken as an input sample and stored at –20°C. Before immunoprecipitation, Dynabeads were washed in 0.1% BSA in PBS and pre-coated with the corresponding primary antibody for at least 3 hours. Protein G Dynabeads (ThermoFisher Scientific) were used to bind mouse immunoglobulins, and Protein A Dynabeads (ThermoFisher Scientific) were used for rabbit immunoglobulins. Chromatin fractions were pre-cleared with washed beads for at least 1 hour at 4°C, magnetic beads were discarded, and chromatin fractions were added to antibody-bound beads, followed by incubation at 4°C overnight, with rotation. The next day, tubes were spun down briefly, supernatant removed, and beads washed for 5 minutes at 4°C with rotation in the following buffers: i. Wash Buffer A (140 mM NaCl, 50 mM HEPES pH 7.9, 1 mM EDTA pH 8, 1% (v/v) Triton X-100, 0.1% SDS (w/v), 0.1% (w/v) Sodium deoxycholate) ii. Wash Buffer B (500 mM NaCl, 50 mM HEPES pH 7.9, 1 mM EDTA pH 8, 1% (v/v) Triton X-100, 0.1% SDS (w/v), 0.1% (w/v) Sodium deoxycholate), iii. Wash Buffer C (250 mM LiCl, 20 mM Tris-HCl pH 7.5, 1 mM EDTA pH 8, 0.5% (v/v) NP40, 0.5% (w/v) Sodium deoxycholate), iv. TE Buffer (10 mM Tris-HCl pH 7.5, 1 mM EDTA pH 8), v. TE Buffer (second wash). Beads were resuspended in Elution Buffer (10 mM Tris-HCl pH 7.5, 1 mM EDTA pH 8, 1% (v/v) SDS), vortexed briefly and incubated at 65°C for 5 minutes. For both input and eluates from ChIP samples, NaCl (160 mM) and RNase A (20 μg/mL) were added, and tubes were incubated at 65°C, 1000 rpm overnight to reverse crosslinks and digest contaminating RNA. After equilibrating samples at room temperature, EDTA (5 mM) and Proteinase K (200 μg/mL) were added, and samples incubated at 45°C for 2 hours to digest proteins. DNA from ChIP and input samples were purified using a ChIP DNA Clean & Concentrator Kit (Zymo) following the manufacturer’s instructions and DNA libraries were prepared using the NEBNext Ultra II DNA Library Prep Kit for Illumina (NEB). Library quality was assessed using a Bioanalyzer High-Sensitivity DNA Analysis kit (Agilent). Samples were then quantified, pooled, and subjected to high-throughput sequencing. Sequencing was performed in a 2×65 bp pair read format using an Illumina NextSeq 2000 flow cell at the MRC Laboratory of Molecular Biology.

### Assay for transposase-accessible chromatin with sequencing (ATAC-seq)

ATAC-seq was performed following the protocol outlined by (Buenrostro et al., 2015). Cells were harvested enzymatically (StemPro Accutase, Gibco) to obtain a single-cell suspension, washed in ice-cold DPBS and counted. 50,000 cells were lysed in cold lysis buffer (10 mM Tris-HCl, pH 7.4, 10 mM NaCl, 3 mM MgCl2 and 0.1% NP-40) to obtain nuclei extracts. Nuclei were immediately transferred to the transposase reaction (25 μL 2× TD buffer, 2.5 μL Tn5 transposase (Illumina) and 22.5 μL nuclease-free water) for 30 minutes at 37 °C. After DNA purification (MinElute PCR Purification Kit, QIAGEN), library fragments were amplified and indexed using the KAPA HiFi HotStart ReadyMix (Roche) and custom Nextera PCR primers as previously described (Buenrostro et al., 2013). Libraries were purified using SPRIselect beads (Beckman Coulter), and their quality assessed using a Bioanalyzer High-Sensitivity DNA Analysis kit (Agilent). Finally, libraries were quantified, pooled, and sequenced in a 2×150 bp pair read format using an Illumina NovaSeq 6000 flow cell at the CRUK Genomics Core Facility.

### CUT&RUN

CUT&RUN (Meers et al., 2019) was performed using the CUT&RUN Assay Kit (Cell Signaling Technology). In summary, 100,000 cells were harvested using StemPro Accutase (Gibco), washed, and then immobilized onto activated Concanavalin A-coated magnetic beads before permeabilisation in a digitonin-containing buffer. The bead-cell complex was then subjected to overnight incubation with the corresponding antibody in a 100 μL volume at 4°C. Following washes, the cells were resuspended in a 50 μL pAG/Mnase mix and further incubated for 1 hour at 4°C. The pAG-Mnase was activated by adding ice-cold calcium-containing buffer and samples were incubated at 4°C for 30 minutes before stopping the reaction. DNA fragments were released into the solution by incubating the beads at 37°C for 30 minutes, followed by separation on a magnet stand, and storage of the supernatant at –20°C. DNA purification from both input and enriched chromatin samples was performed using DNA Purification Buffers and Spin Columns for ChIP and CUT&RUN (Cell Signaling Technology). Subsequently, CUT&RUN libraries were constructed using the NEBNext Ultra II DNA Library Prep Kit for Illumina (NEB) with the assistance of a Biomek FXP Laboratory Automation Workstation (Beckman Coulter), with minor modifications to preserve small DNA fragments as recommended. The quality of the libraries was assessed using a Bioanalyzer High-Sensitivity DNA Analysis kit (Agilent), followed by quantification, pooling, and high-throughput sequencing. Sequencing was performed in a 2×65 bp pair read format using an Illumina NextSeq 2000 flow cell at the MRC Laboratory of Molecular Biology. Positive (H3K4me3) and negative (Rabbit IgG) controls were performed in parallel.

### bTMP-seq

Biotinylated 4,5,8-trimethylpsoralen (bTMP) was synthesised as previously described (Saffran *et al*., 1988) and purified by flash column chromatography on silica. The purity and structure of the compound were confirmed by mass and 1H NMR spectroscopy. The powder was protected from light, and 1 mg/mL stocks were dissolved in DMSO, stored at –20°C, and protected from light until and during use.

Biotin TMP-seq was performed in duplicate as previously described (Naughton *et al*., 2013; Achar *et al*., 2020) with modifications to adapt the method to high-throughput sequencing. 1 – 2 million cells were incubated with bTMP (at 50 μg/mL or 150 μg/mL) in 1ml final volume in darkness for 20 minutes at 37°C. The psoralen was then crosslinked to DNA with 800 mJ.cm^-2^ 365 nm UV light. Following two washes with PBS, cells were harvested with Accutase and the pellets snap frozen until all conditions were collected. Pellets were resuspended in 100 µl cold PBS and DNA was extracted using the Monarch Genomic DNA Purification Kit (NEB) with RNase A and Proteinase K digestion for 2 hours. The cross-linked DNA was sheared to an average size of 500 bp using a Covaris M220 Ultra Sonicator (Peak Incident Power = 75 W; Duty Factor = 10%; Cycles Per burst = 200; Time = 64 seconds) in microTUBE-50 AFA Fiber Screw-Cap tubes (Covaris). Sonication size was confirmed on a TapeStation (Agilent). Subsequently, bTMP-DNA complexes were captured using Dynabeads MyOne Streptavidin C1 magnetic beads (Invitrogen) and incubated for 3 hours at room temperature with rotation. The beads were washed sequentially for 5 minutes at room temperature with buffers: Wash Buffer I (20 mM Tris-HCl pH 8, 2 mM EDTA, 150 mM NaCl, 1% Triton X, 0.1% SDS), Wash Buffer II (20 mM Tris-HCl pH 8, 2 mM EDTA, 500 mM NaCl, 1% Triton X, 0.1% SDS), Wash Buffer III (250 mM LiCl, 10 mM Tris pH 8.0, 0.5% Na-deoxycholate, 0.5% NP-40, 1 mM EDTA) and twice with 1× TE Buffer (20 mM Tris pH 8.0, 2 mM EDTA). The bTMP-DNA complexes were eluted from the beads in 50 μL elution buffer (95% formamide, 10 mM EDTA) at 65°C for 5 minutes as per manufacturer’s instructions and the eluted samples were purified with a MinElute PCR purification kit (QIAGEN). Following elution, the samples were further sonicated to a target size of 150 bp using a Covaris M220 Ultra Sonicator (Peak Incident Power = 75 W; Duty Factor = 10%; Cycles Per burst = 200; Time = 64 seconds). This reduces the chance of the library templates containing a bTMP molecule that may inhibit amplification, while retaining localisation to the initially enriched 500bp sections. Input DNA from bTMP-treated cells was separated before streptavidin pulldown step.

For the naked DNA control, genomic DNA was extracted from untreated cells using the Monarch Genomic DNA Purification Kit (NEB) and sheared to an average size of 500 bp using a Covaris M220 Ultra Sonicator with the same parameters as above. Biotin TMP was added to purified DNA (50 μg/mL or 150 μg/mL), incubated in the dark for 20 minutes and cross-linked with 800 mJ.cm^-2^ 365 nm UV light. DNA was precipitated using isopropanol, washed with 70% ethanol and dissolved in buffer (50 mM Tris pH 8.0, 10 mM EDTA, 0.1% SDS). The DNA was then incubated with Dynabeads MyOne Streptavidin for 1 hour at room temperature, followed by washing, elution and second sonication to 150 bp, as described above.

DNA libraries were generated using the NEBNext Ultra II DNA Library Prep Kit for Illumina (NEB) and barcoded with NEBNext Multiplex Oligos for Illumina (NEB). Final libraries were quality tested for primer and adapter contamination using an Agilent 2100 Bioanalyzer High Sensitivity DNA chip. The samples were quantified, pooled, and subjected to high-throughput sequencing using an Illumina Nextseq 2000 instrument.

### Hi-C chromosome conformation capture

Hi-C analysis was performed on hESCs using the Arima-Hi-C kit (Arima Genomics). Initially, cells were harvested with StemPro Accutase, washed in PBS and crosslinked in 2% formaldehyde solution (Sigma-Aldrich) at room temperature for 10 minutes before quenching the reaction. Crosslinked cells were then aliquoted, pelleted (500 g, 5 minutes), snap-frozen in liquid nitrogen, and preserved at –80°C. For each Arima-Hi-C reaction, crosslinked cells corresponding to a minimum of 5 μg of DNA were thawed, lysed, and subjected to chromatin digestion using a restriction enzyme cocktail. Subsequently, newly generated overhangs were labelled with biotinylated nucleotides, and spatially proximal digested DNA ends were ligated. Before library preparation, DNA was fragmented to 400 bp using a Covaris M220 Ultra Sonicator (Peak Incident Power = 50 W; Duty Factor = 10%; Cycles Per burst = 200; Time = 125 seconds) and microTUBE AFA Fiber Pre-Slit Snap-Cap 6×16mm tubes (Covaris). Following size selection with AMPure XP beads (Beckman Coulter), biotinylated fragments were enriched. Swift Biosciences Accel-NGS 2S Plus DNA Library Kit and Swift Biosciences Indexing Kit reagents were employed for end-repair and adapter ligation, following the modified protocol provided with the Arima-Hi-C Kit. The bead-bound Hi-C DNA was then subjected to 6-8 cycles of amplification using KAPA Library Amplification Kit (Roche) reagents. Final libraries were tested for primer and adapter contamination using an Agilent 4150 TapeStation and D500 ScreenTape Assay Reagent Kit (Agilent). Finally, libraries were quantified, pooled, and prepared for high-throughput sequencing. Sequencing was performed in a 2×150 bp pair read format using an Illumina NextSeq 2000 flow cell at the MRC Laboratory of Molecular Biology.

### Human reference genome and annotation

All analyses were performed on the hg38/GRCh38 human reference genome assembly. Datasets originally obtained with coordinates on other assemblies were projected onto the hg38 assembly using the liftOver function from the rtracklayer R/Bioconductor package (Lawrence *et al*., 2009), with the corresponding chain files obtained from http://hgdownload.cse.ucsc.edu in the individual assembly download sections. For RNAseq Ensembl gene annotation GRCh38.87 was used, while for other analyses annotations were extracted using the biomaRt R/Bioconductor package (Durinck *et al*., 2009). We considered only protein coding genes (n = 19,254). Enhancers were defined, from ChIP-seq and ATAC-seq datasets, by using H3K4me1 peaks without H3K4me3 but overlapping ATAC-seq Tn5 hypersensitive site peaks. Enhancers overlapping TSSs (TSS ± 1 kb) were excluded. Among these enhancers, those overlapping H3K27ac peaks were taken as active enhancers, those overlapping with H3K27me3 peaks were taken as poised enhancers, those overlapping with neither H3K27ac nor H3K27me3 peaks were considered primed enhancers. Stitched enhancers (or superenhancer domains) were defined according to Whyte et al. (Whyte *et al*., 2013) by merging enhancer domains separated by less than 12.5 kb (n = 3,019).

### Initial bioinformatics data processing

For RNAseq, ATACseq, ChIPseq and CUT&RUN analysis, sequencing reads were trimmed for quality with Trim Galore (doi: 10.5281/zenodo.5127898), with a minimum quality score of 30. For bTMP-seq, ATACseq, ChIPseq and CUT&RUN analysis reads were aligned to the hg38/GRCh38 human reference genome assembly using bowtie2 (Langmead and Salzberg, 2012), with the following parameters: bTMP-seq parameters: default; ATACseq parameters: –-local; ChIPseq parameters: –-local; CUT&RUN parameters: –-local –-very-sensitive-local –-no-unal –-no-mixed –-no-discordant –I 10 –X 700. For RNAseq analysis reads were aligned using tophat2 (Kim *et al*., 2013) with default parameters, and then counted using htseq-count (Anders *et al*., 2015). For bTMP-seq, ATACseq and CUT&RUN analysis non-uniquely mapped reads were discarded. Aligned reads were then sorted and PCR duplicates were filtered out with samtools (Li *et al*., 2009). For ChIPseq PCR duplicates were filtered out with Picard MarkDuplicates (Institute, 2019). For RNAseq, bTMP-seq, ATACseq, ChIPseq and CUT&RUN analysis read counts were then computed at 50 bp and 50 kb resolutions using the bamCoverage function of deeptools (Ramírez *et al*., 2016) using the following parameters: –-smoothLength 350 –-extendReads 150 and –-smoothLength 150000 –-extendReads 150.

### RNAseq analysis

Raw gene-level count files from three biological replicates per condition (UD, H24, H48, H72) were analysed with edgeR (Robinson *et al*., 2010): libraries were created with DGEList, normalised using trimmed mean of M-values (TMM; calcNormFactors), and converted to RPKM (rpkm) using gene lengths (Robinson and Oshlack, 2010). Replicate RPKMs were averaged within each condition, and values <0.01 were set to 0. Duplicate gene IDs were aggregated by summation, lowly expressed genes were filtered with filterByExpr, and log2 counts per million (logCPM) were computed (cpm, log=TRUE, prior.count=1). The top 500 most variable genes were selected by variance across samples and displayed as a heatmap using gplots heatmap.2 (https://CRAN.R-project.org/package=gplots) with row-wise Z-scaling, fixed z-limits (−3 to 3). Samples were ordered by condition (UD, H24, H48, H72) and replicate, rows were hierarchically clustered. GO terms were extracted with the API from g:Profiler (Reimand *et al*., 2007).

### bTMP-seq analysis

Initial alignment was carried out as above, with read counts computed in 50bp and 50kb bins. Bin read counts were normalised in Counts Per Million (CPM) mapped reads. After assessing reproducibility, by comparing normalised read counts for the immunoprecipitated samples, duplicates were merged. Supercoiling scores were computed for each genomic window using bTMP IP and input read counts. Read counts for bTMP binding ‘in cells’ (log2(IP/input)) was subtracted from the ‘naked genomic DNA’ score (log2(IP/input)) to correct for false positive binding of bTMP.

### ATACseq, CUT&RUN and ChIP-seq analysis

Initial alignment was carried out as above, with read counts computed in 50bp bins. For both CUT&RUN and ChIP-seq counts were log2 normalised to their respective inputs with the bigwigCompare function of deepTools (Ramírez *et al*., 2016). The resulting bigwig files were used for subsequent analysis. After assessing reproducibility, duplicates were merged. Sites of RNA Pol II, RAD21 and CTCF accumulation were identified by peak calling performed with the callpeak function of MACS2 (Gaspar, 2018) using the narrow mode. Only the top 10,000 peaks were considered for the analysis. ChIP-seq peaks for histone modifications and ATAC-seq were called with the bdgpeakcall function of MACS2 (Gaspar, 2018), using the following cut-off and maximum gap parameters, ATAC-seq (−c 0.1 –g 50), H3K4me1 (−c 0.3 –g 50), H3K4me3 (−c 0.3 –g 50), H3K27ac (−c 1.3 –g 50) and H3K27me3 (−c 0.3 –g 50).

### Pol II occupancy and eRNA expression

RNA Pol II occupancy within genes and eRNA expression at enhancers were computed from Pol II ChIP-seq and RNA-seq data respectively. Pol II occupancy within protein coding genes was computed by considering domains starting 1 kb after TSSs and finishing 1 kb before TTSs to exclude sites known for pausing of non-transcriptionally active Pol II. Only genes longer than 3kb were considered for the analysis. Pol II ChIP-seq and RNA-seq read counts were computed in 50 bp windows using the bamCoverage function of deeptools. Pol II occupancy was defined as the sum of reads overlapping gene domains and normalising for their lengths. eRNA expression was computed as Reads Per Kilobase per Million (RPKM) mapped reads considering RNA-seq reads overlapping stitched enhancer domains.

### Metagene analysis

Averages of supercoiling, ChIP-seq, CUT&RUN and other features over protein coding genes, topologically associating domains (TADs) and A/B compartments were computed using the normalizeToMatrix function of the EnrichedHeatmap R/Bioconductor package (Gu *et al*., 2018). Protein coding genes and their adjacent domains (± 10 kb) were divided into 100 equally sized windows and signal averages were computed using aggregated data at 50 nt resolution. TADs and their adjacent domains (± 500 kb) were divided into 1000 equally sized windows and signal averages were computed using aggregated data at 50 kb resolution. A/B compartments and their adjacent domains (± 2 Mb) were divided into 1000 equally sized windows and signal averages were computed using aggregated data at 50 kb resolution. Genomic signals were normalised to target regions using the coverage mode of normalizeToMatrix omitting bins with missing values in the computation. Enrichment values were then computed by averaging genomic signals across windows within regions of interest and corrected for background values computed from adjacent domains.

### Hi-C data processing

Hi-C reads were aligned to the hg38/GRCh38 human reference genome assembly using BWA-MEM (Li, 2013) with the following parameters, –A1 –B4 –E50 –L0. Aligned reads were then processed and analysed with Hi-CExplorer (Ramírez *et al*., 2018). Genomic locations of restriction sites were identified using the hicFindRestSite function and the ‘GATC’ and ‘GANTC’ patterns. Raw cis contact matrices were computed using the hicBuildMatrix function at 1 kb resolution. All experiments were performed in duplicate. After assessing the quality of the generated matrices, replicates were merged using the hicSumMatrices function. Contact matrices were balanced using the hicCorrectMatrix function and the Iterative Correction and Eigenvector decomposition (ICE) method (Imakaev *et al*., 2012). The resulting matrices in .h5 format were converted into the .cool format using the hicConvertFormat if required.

### A/B compartment identification

Genome-wide A/B compartment status was evaluated using the hicPCA function from ICE balanced Hi-C matrices for each chromosome and the ‘Lieberman’ method (Lieberman-Aiden *et al*., 2009). Coordinates corresponding to transitions between positive and negative eigenvector values demarcate boundaries of compartments. Domains separated by distances smaller than 1 Mb were merged. We assigned positive eigenvector values to the transcriptional active A compartment, and negative values to the transcriptional inactive B compartment. Transcriptional activity was determined based on our RNA-seq data. Chromosomes showing poor compartmentalisation in either the undifferentiated or definitive endoderm states, i.e. chr16, chr21 and chr22, were excluded from the analysis. Intervals overlapping with the ENCODE phase 4 exclusion list regions, available at https://www.encodeproject.org/files/ENCFF356LFX/, were also excluded.

### Topologically associating domains

Topologically associating domains (TADs) were identified using the hicFindTADs function from ICE balanced Hi-C matrices at 25 and 250 kb resolutions. The parameters used were –-step 25000 –-minDepth 150000 –-maxDepth 300000 –-minBoundaryDistance 200000 and –-step 250000 –-minDepth 1500000 –-maxDepth 3000000 –-minBoundaryDistance 2000000. TAD scores were corrected for multiple testing using the false discovery rate (FDR) method and only TADs with q-values above 0.01 were considered. Identified TADs were then assigned to A/B compartments based a minimum overlap distance of 50 kb with the relevant domain type. TADs wider than 10 times the maximum window length considered for TADs identification, i.e. 3 or 30 Mb depending on the resolution, were excluded from the analysis. TADs overlapping with the ENCODE phase 4 exclusion list regions were also excluded. Insulation score. Insulation scores (ISs) were computed using a diamond-window score approach with the diamond-insulation function of cooltools from ICE the balanced Hi-C matrices at 1, 10, 20 and 50 kb resolutions (Abdennur *et al*., 2022). Diagonals were considered for computation. Tiles displaying values outside the 1 to 99 % quantiles were excluded from the analysis. IS data were converted into bigwig files using the GenomicRanges R/Bioconductor package (Lawrence *et al*., 2013) for downstream analysis.

### Hi-C pileups

Pile-up analysis on Hi-C data was performed using Coolpup.py (Flyamer *et al*., 2020). Hi-C pileups were computed from unbalanced Hi-C matrices with the following parameters, –-rescale –-rescale_pad 1 –-rescale_size 199 –-ignore_diags 0 –-unbalance –-coverage_norm. Normalisation to expected values was performed by dividing the pileups over randomly shifted control regions (−−nshifts 10) according to the Aggregate Peak Analysis (APA) method (Rao *et al*., 2014). Pileups at protein-coding genes were performed using Hi-C matrices at 1 kb resolution and considering only genes longer than 30 kb. Pileups at TADs were performed using Hi-C matrices at 50 kb resolution. Resulting pileups were plotted using a custom R-script.

### Computational and statistical analyses

Analysis and all statistical calculations were performed in R (version 4.5.2). All scripts are available at https://github.com/Sale-lab/ScDiff.

## Acknowledgements & Funding

We would like to thank Guillaume Guilbaud, Linda van Bijsterveldt, Jan Verburg and members of the Sale group for discussions and for critical reading of the manuscript. This work was supported by core MRC funding to LMB (MC_U105178808). C.P. was supported by a César Milstein studentship from the Darwin Trust of Edinburgh. A.Z. was supported by a PhD fellowship from the Boehringer Ingelheim Fonds and a Neuberger Studentship from the Max Perutz Fund in collaboration with Trinity College. J.E.S. thanks the Wellcome Sanger Institute for support through the associate faculty program.

## Data Availability

Raw and processed sequencing data are available at GEO (www.ncbi.nlm.nih.gov/geo/) under accession numbers GSE318996 and GSE202799. All scripts are available at https://github.com/Sale-lab/ScDiff.

## Supplementary Figures

**Figure S1.**
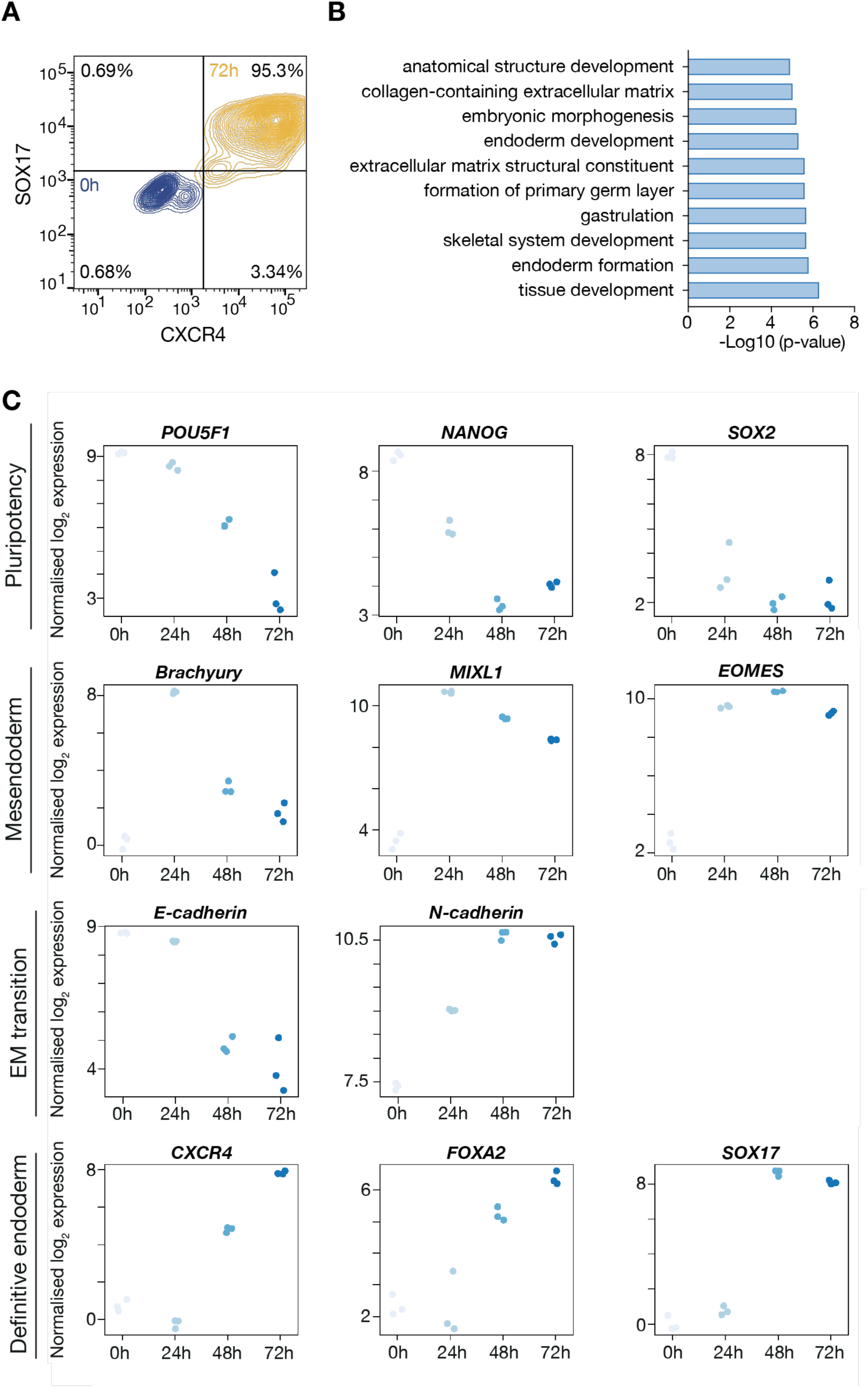
Expression changes in key genes involved in the transition from pluripotency to definitive endoderm (DE) during the 72-hour differentiation protocol used in this study. A. The efficiency of H9 differentiation to DE monitored by permeabilisation-based flow cytometry for SOX17 and CXCR4 expression. B. Most enriched gene ontology (GO) terms in significantly upregulated genes between 0 and 72 hours. p calculated with the hypergeometric test with Benjaminin-Hochberg FDR multiple testing correction. C. RNA-seq expression patterns of key genes involved in DE differentiation in H9 cells transitioning from pluripotency to DE. Each blue dot represents a biological replicate sampled for each time point.

**Figure S2.**
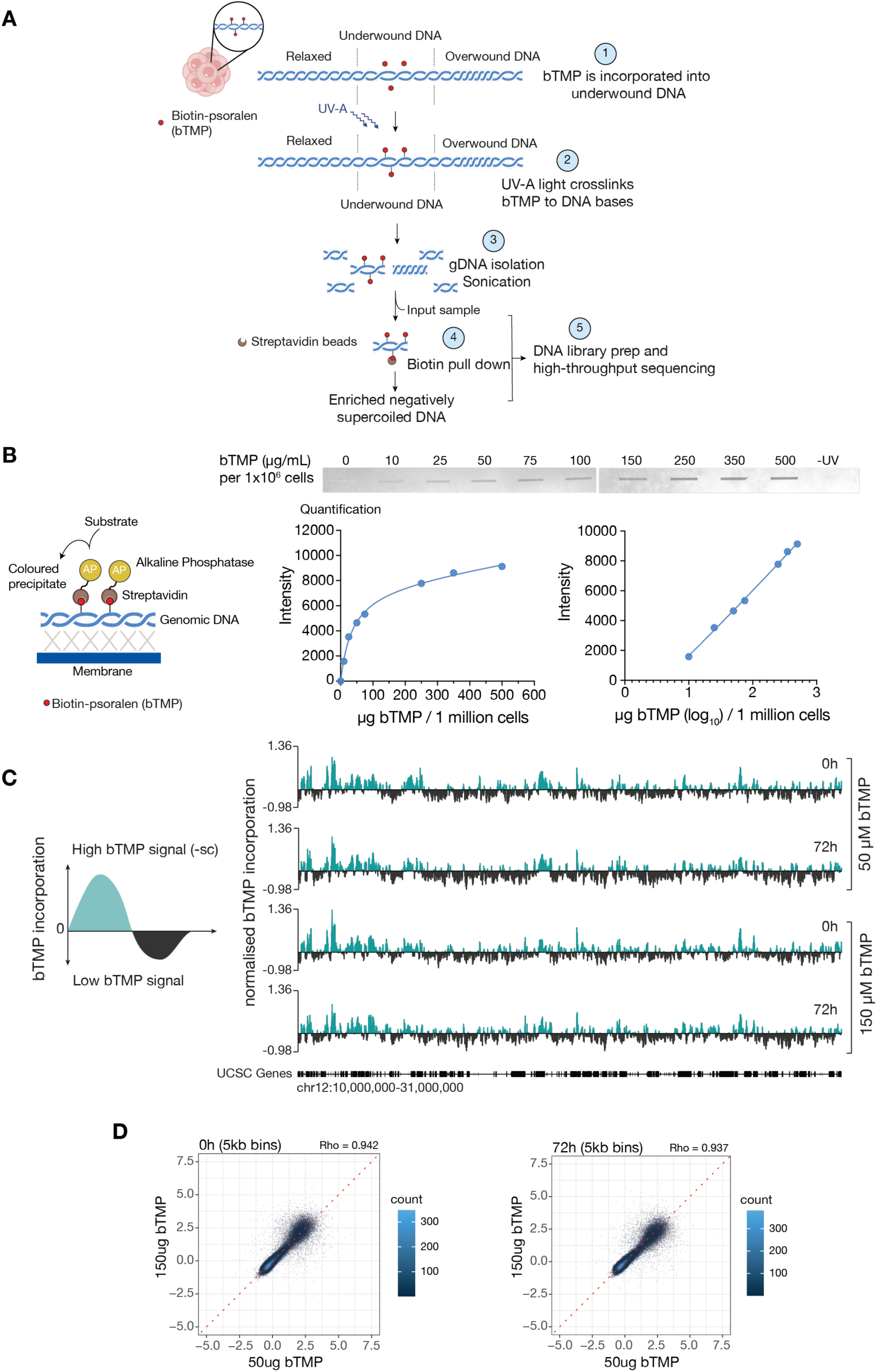
bTMP-seq workflow and controls. A. Schematic summarising the experimental workflow of biotinylated trimethylpsoralen-seq (bTMP-seq) for mapping DNA supercoiling. Following treatment with psoralen for 20 minutes at 37°C in the dark (1), cells are exposed to 360-nm UVA light to crosslink the Biotinylated TMP to the DNA where it has intercalated (2). At the limiting doses used, bTMP intercalation correlates with the degree of negative supercoiling in the DNA (Sinden *et al*., 1980). Genomic DNA is then purified and sonicated (3). Negatively supercoiled DNA is enriched using streptavidin-coated magnetic beads (4), amplified, and sequenced alongside an input control. B. Slot blot assay to measure bTMP incorporation into genomic DNA (0.2 μg per well). Left panel: bTMP is detected with streptavidin-conjugated alkaline phosphatase and a biotin chromogenic detection kit (ThermoFisher). The intensity of the bands was quantified using Image J software, plotted on Graph Prism and non-linear fit curves adjusted using Graph Prism. Individual measurements are shown in blue dots. The right hand plot is a zoomed in version of the left hand plot showing that, at the ratio of bTMP to cells used, incorporation is linear. C. Example IGV tracks of normalised bTMP-seq data over a 21 Mb interval of Chromosome 12. Bin size 50 kb. USCS genes shown as black boxes. D. Comparison of bTMP-seq in H9 cells at t = 0h (left panel) or t = 72h (right panel) treated with either 50 or 150 μg/mL of bTMP. Each dataset is made up of data from three biological replicates. The scatter plots show two-way Pearson correlations of the supercoiling score (normalised to input only) between different doses of bTMP. All experiments reported in this paper were performed with 50 μg/mL bTMP.

**Figure S3.**
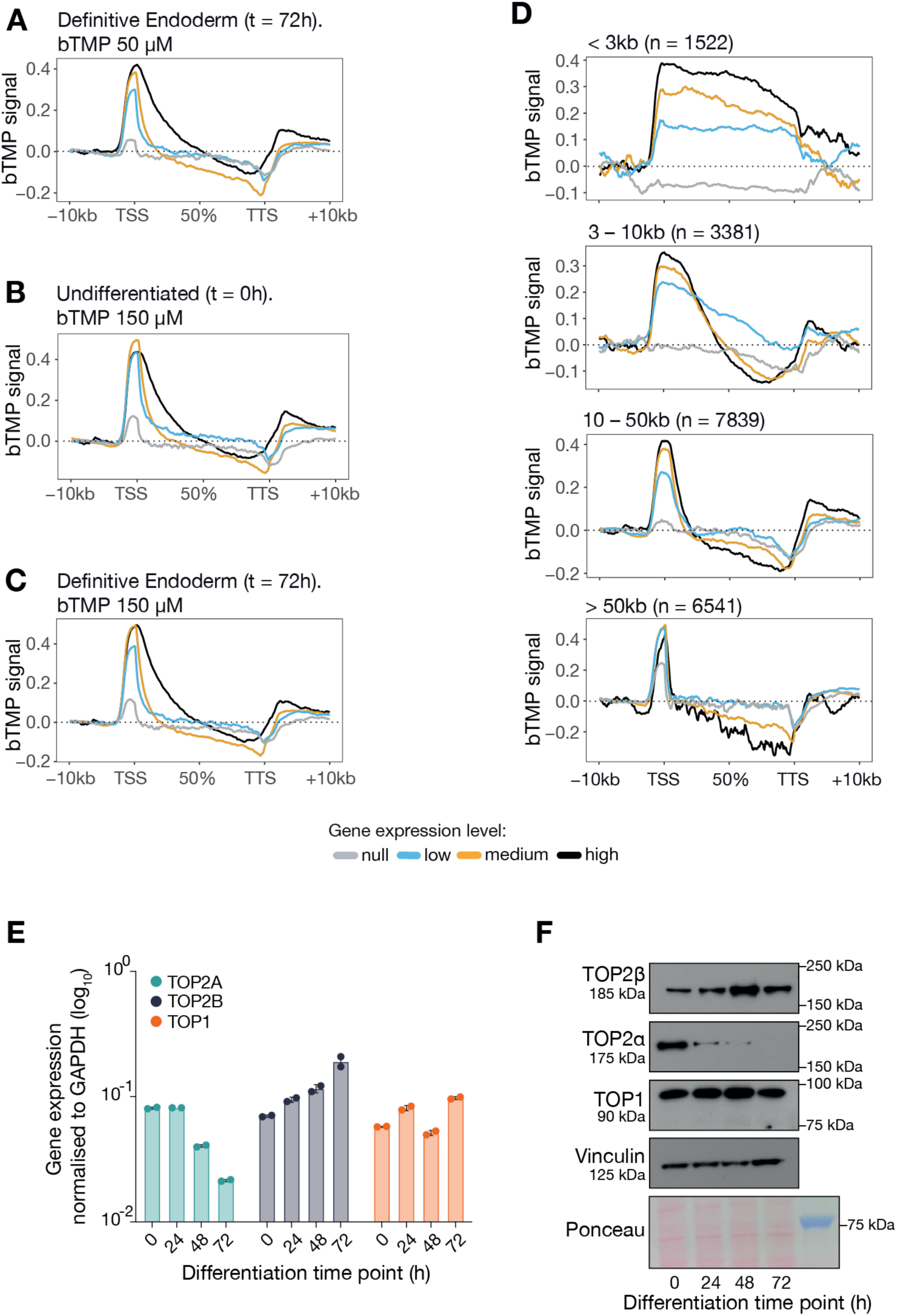
A-D. Metagenes of normalised bTMP incorporation in H9 cells at various concentrations and timepoints during differentiation. The expression groups are no expression – light grey; low (quantile < 0.5), blue; medium (quantile = 0.5 – 0.9), orange; and high (quantile > 0.9), dark grey. Coverage matrices were computed in 500 bp windows. Mean changes (enrichment over background) were computed in 400 bins (100 in each flank, 200 in gene) and normalised using background values from domains adjacent to genes (−10 to –5 kb). TSS = transcription start site; TTS = transcription termination site. A. bTMP = 50 µM (as in Figure 2B) in H9 cells differentiated to definitive endoderm (t = 72h). B. bTMP = 150 µM in undifferentiated H9 cells (t = 0h). C. bTMP = 150 µM in H9 cells differentiated to definitive endoderm (t = 72h). D. bTMP = 50 µM (as in Figure 2B) in undifferentiated H9 cells (t = 0h). Genes are divided by total length. E. Expression of *TOP2A*, *TOP2B* and *TOP1* monitored by real-time qPCR in undifferentiated H9 cells and through endoderm differentiation (24-72h); mean±SD of at two biological replicates (n=2) averaged across three technical replicates are plotted. F. Immunoblot of TOP1, TOP2α and TOP2β in H9 cells differentiating to definitive endoderm. Samples were collected every 24 hours. Loading controls: vinculin and Ponceau S staining of the membrate. Size markers: Precision Plus Protein All Blue Prestained Protein Standards (Bio-Rad).

**Figure S4.**
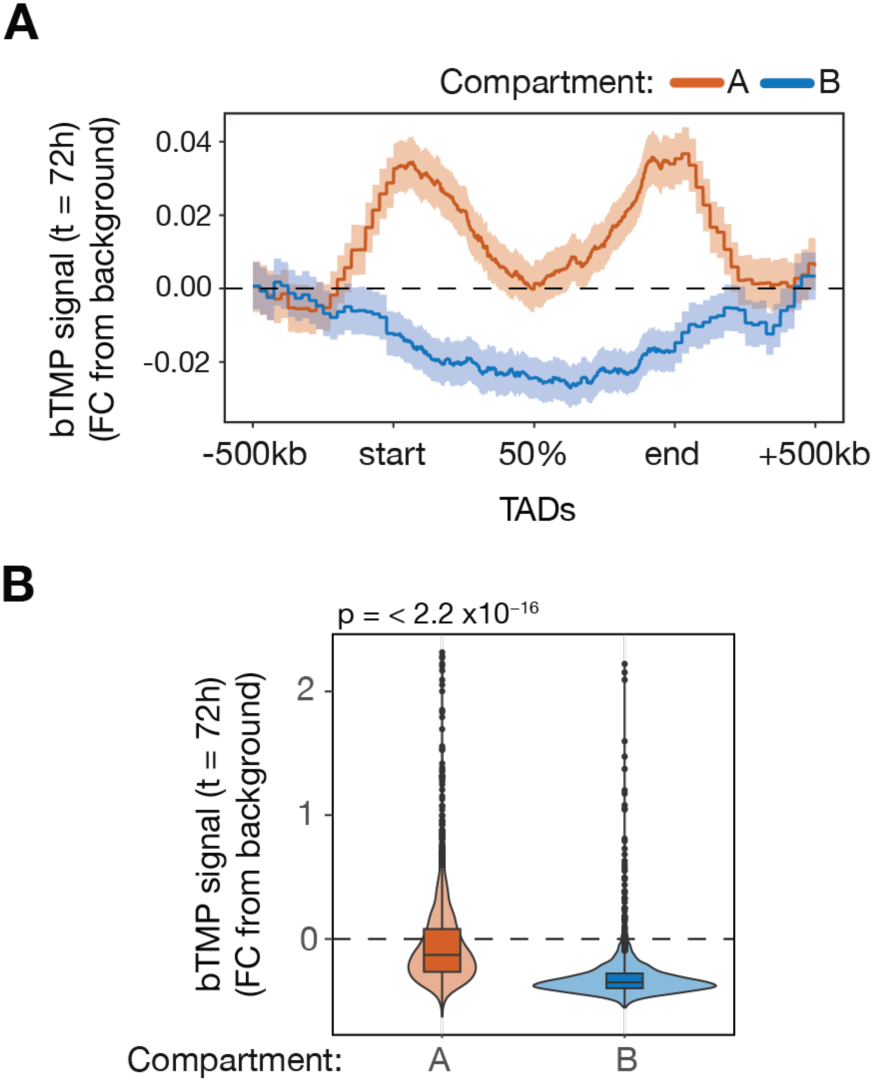
A. bTMP-seq signal variation normalised to background within TADs at t = 72h binned by compartment (A = red; B = blue). TAD boundaries are denoted as ‘start’ and ‘end’ and the middle meta-distance is denoted as 50%. B. Quantitation of bTMP-seq signal in TADs of the A (red) and B (blue) compartments at t = 72h. P value calculated with a two-sample Kolmogorov-Smirnov test.

**Figure S5.**
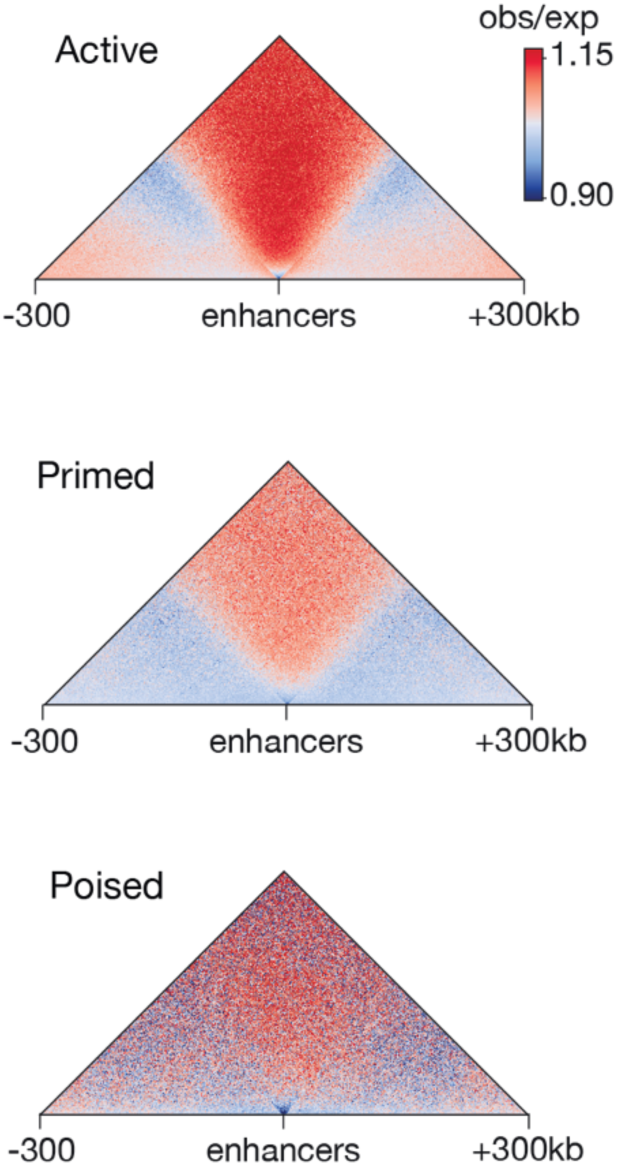
Hi-C contact maps centred on enhancers +/− 300 kb categorised as Active, Primed and Poised. See Methods.

**Table S1.** The top 500 most variable genes between 0 h (pluripotency) and 72 h (definitive endoderm). See separate .csv file: ‘Table S1_top500_most_variable_genes.csv’

**Table S2.** Table of antibodies used in this study.

| Target | Source | Identifier | Species | Type | Conjugated | Dilution | Use |
| --- | --- | --- | --- | --- | --- | --- | --- |
| SOX17 | BD Pharmingen | 562205 | Mouse IgG1, $\kappa$ | Monoclonal | Alexa Fluor 488 | 1:20 | Flow cytometry |
| EOMES | Invitrogen | 50-4877-42 | Mouse IgG1, $\kappa$ | Monoclonal | eFluor660 | 1:20 | Flow cytometry |
| CXCR4 (CD184) | BD Pharmingen | 557145 | Mouse IgG2a, $\kappa$ | Monoclonal | PE | 1:20 | Flow cytometry |
| Alexa Fluor 488 Mouse IgG1, $\kappa$ Isotype Control | BD Pharmingen | 557782 | Mouse IgG1 $\kappa$ | Monoclonal | Alexa Fluor 488 | 1:20 | Flow cytometry |
| eFluor 660 Mouse IgG1 $\kappa$ Isotype Control | Invitrogen | 50-4714-82 | Mouse IgG1 $\kappa$ | Monoclonal | eFluor 660 | 1:20 | Flow cytometry |
| PE Mouse IgG2a, $\kappa$ Isotype Control | BD Pharmingen | 556653 | Mouse IgG2a $\kappa$ | Monoclonal | PE | 1:20 | Flow cytometry |
| CUT & RUN Isotype Control | Cell Signaling | #3900 | Rabbit IgG | Monoclonal | - | 1:100 | CUT&RUN |
| H3K4me3 | Cell Signaling | #9751 | Rabbit | Monoclonal | - | 1:50 | CUT&RUN |
| H3K27me3 | Cell Signaling | #9733 | Rabbit | Monoclonal | - | 1:50 | CUT&RUN |
| H3K4me1 | Abcam | ab8895 | Rabbit | Polyclonal | - | 1:50 | CUT&RUN |
| H3 | Abcam | ab1791 | Rabbit | Polyclonal | - | 1:50 | CUT&RUN |
| CTCF | Cell Signaling | #3418 | Rabbit | Monoclonal | - | 1:50 | CUT&RUN |
| TOP1 | Abcam | ab3825 | Rabbit | Polyclonal | - | 1:50 | CUT&RUN |
| TOP2β | Abcam | ab58442 | Rabbit | | - | 1:50 | CUT&RUN |
| TOP2α | Abcam | ab52934 | Rabbit | Monoclonal | - | 1:50 | CUT&RUN |
| RAD21 | Abcam | ab992 | Rabbit | Polyclonal | - | 6μg | ChIP-seq |
| RPB1 CTD (4H8) | Cell Signaling | #2629 | Mouse | Monoclonal | - | 8μg | ChIP-seq |

## References

1. Abdennur N, Abraham S, Fudenberg G, Flyamer IM, Galitsyna AA, Goloborodko A, Imakaev M, Oksuz BA, Venev SV (2022) Cooltools: enabling high-resolution Hi-C analysis in Python. bioRxiv, 40

2. Achar YJ, Adhil M, Choudhary R, Gilbert N, Foiani M (2020) Negative supercoil at gene boundaries modulates gene topology. Nature, 577: 701–705

3. Anders S, Pyl PT, Huber W (2015) HTSeq--a Python framework to work with high-throughput sequencing data. Bioinformatics, 31: 166–169

4. Andersson R, Sandelin A (2020) Determinants of enhancer and promoter activities of regulatory elements. Nat Rev Genet, 21: 71–87

5. Baranello L, Wojtowicz D, Cui K, Devaiah BN, Chung HJ, Chan-Salis KY, Guha R, Wilson K, Zhang X, Zhang H, Piotrowski J, Thomas CJ, Singer DS, Pugh BF, Pommier Y, Przytycka TM, Kouzine F, Lewis BA, Zhao K, Levens D (2016) RNA Polymerase II Regulates Topoisomerase 1 Activity to Favor Efficient Transcription. Cell, 165: 357–371

6. Beagrie RA, Scialdone A, Schueler M, Kraemer DC, Chotalia M, Xie SQ, Barbieri M, de Santiago I, Lavitas LM, Branco MR, Fraser J, Dostie J, Game L, Dillon N, Edwards PA, Nicodemi M, Pombo A (2017) Complex multi-enhancer contacts captured by genome architecture mapping. Nature, 543: 519–524

7. Benedetti F, Dorier J, Burnier Y, Stasiak A (2014) Models that include supercoiling of topological domains reproduce several known features of interphase chromosomes. Nucleic Acids Res, 42: 2848–2855

8. Benedetti F, Racko D, Dorier J, Burnier Y, Stasiak A (2017) Transcription-induced supercoiling explains formation of self-interacting chromatin domains in S. pombe. Nucleic Acids Res, 45: 9850–9859

9. Bermúdez I, García-Martínez J, Pérez-Ortín JE, Roca J (2010) A method for genome-wide analysis of DNA helical tension by means of psoralen-DNA photobinding. Nucleic Acids Res, 38: e182

10. Björkegren C, Baranello L (2018) DNA Supercoiling, Topoisomerases, and Cohesin: Partners in Regulating Chromatin Architecture. Int J Mol Sci, 19: E884

11. Bonev B, Cavalli G (2016) Organization and function of the 3D genome. Nat Rev Genet, 17: 661–678

12. Institute B (2019) Picard Toolkit.

13. Busslinger GA, Stocsits RR, van der Lelij P, Axelsson E, Tedeschi A, Galjart N, Peters JM (2017) Cohesin is positioned in mammalian genomes by transcription, CTCF and Wapl. Nature, 544: 503–507

14. Canela A, Maman Y, Huang SN, Wutz G, Tang W, Zagnoli-Vieira G, Callen E, Wong N, Day A, Peters JM, Caldecott KW, Pommier Y, Nussenzweig A (2019) Topoisomerase II-Induced Chromosome Breakage and Translocation Is Determined by Chromosome Architecture and Transcriptional Activity. Mol Cell, 75: 252–266.e8

15. Crane E, Bian Q, McCord RP, Lajoie BR, Wheeler BS, Ralston EJ, Uzawa S, Dekker J, Meyer BJ (2015) Condensin-driven remodelling of X chromosome topology during dosage compensation. Nature, 523: 240–244

16. Davidson IF, Barth R, Zaczek M, van der Torre J, Tang W, Nagasaka K, Janissen R, Kerssemakers J, Wutz G, Dekker C, Peters JM (2023) CTCF is a DNA-tension-dependent barrier to cohesin-mediated loop extrusion. Nature, 616: 822–827

17. Deng W, Lee J, Wang H, Miller J, Reik A, Gregory PD, Dean A, Blobel GA (2012) Controlling long-range genomic interactions at a native locus by targeted tethering of a looping factor. Cell, 149: 1233–1244

18. Dixon JR, Selvaraj S, Yue F, Kim A, Li Y, Shen Y, Hu M, Liu JS, Ren B (2012) Topological domains in mammalian genomes identified by analysis of chromatin interactions. Nature, 485: 376–380

19. Durinck S, Spellman PT, Birney E, Huber W (2009) Mapping identifiers for the integration of genomic datasets with the R/Bioconductor package biomaRt. Nat Protoc, 4: 1184–1191

20. Eldridge CB, Allen FJ, Crisp A, Grandy RA, Vallier L, Sale JE (2020) A p53-Dependent Checkpoint Induced upon DNA Damage Alters Cell Fate during hiPSC Differentiation. Stem Cell Reports, 15: 827–835

21. Fernández X, Díaz-Ingelmo O, Martínez-García B, Roca J (2014) Chromatin regulates DNA torsional energy via topoisomerase II-mediated relaxation of positive supercoils. EMBO J, 33: 1492–1501

22. Fisher JB, Pulakanti K, Rao S, Duncan SA (2017) GATA6 is essential for endoderm formation from human pluripotent stem cells. Biol Open, 6: 1084–1095

23. Flyamer IM, Illingworth RS, Bickmore WA (2020) Coolpup.py: versatile pile-up analysis of Hi-C data. Bioinformatics, 36: 2980–2985

24. Fogg JM, Judge AK, Stricker E, Chan HL, Zechiedrich L (2021) Supercoiling and looping promote DNA base accessibility and coordination among distant sites. Nat Commun, 12: 5683

25. Fortin JP, Hansen KD (2015) Reconstructing A/B compartments as revealed by Hi-C using long-range correlations in epigenetic data. Genome Biol, 16: 180

26. Freire-Pritchett P, Schoenfelder S, Várnai C, Wingett SW, Cairns J, Collier AJ, García-Vílchez R, Furlan-Magaril M, Osborne CS, Fraser P, Rugg-Gunn PJ, Spivakov M (2017) Global reorganisation of cis-regulatory units upon lineage commitment of human embryonic stem cells. Elife, 6: e21926

27. Fu Z, Guo MS, Zhou W, Xiao J (2024) Differential roles of positive and negative supercoiling in organizing the E. coli genome. Nucleic Acids Res, 52: 724–737

28. Gaspar JM (2018) Improved peak-calling with MACS2. bioRxiv, 126

29. Gilbert N, Marenduzzo D (2025) Topological epigenetics: The biophysics of DNA supercoiling and its relation to transcription and genome instability. Curr Opin Cell Biol, 92: 102448

30. Gu Z, Eils R, Schlesner M, Ishaque N (2018) EnrichedHeatmap: an R/Bioconductor package for comprehensive visualization of genomic signal associations. BMC Genomics, 19: 234

31. Guo MS, Kawamura R, Littlehale ML, Marko JF, Laub MT (2021) High-resolution, genome-wide mapping of positive supercoiling in chromosomes. Elife, 10: e67236

32. Haarhuis JHI, van der Weide RH, Blomen VA, Yáñez-Cuna JO, Amendola M, van Ruiten MS, Krijger PHL, Teunissen H, Medema RH, van Steensel B, Brummelkamp TR, de Wit E, Rowland BD (2017) The Cohesin Release Factor WAPL Restricts Chromatin Loop Extension. Cell, 169: 693–707.e14

33. Imakaev M, Fudenberg G, McCord RP, Naumova N, Goloborodko A, Lajoie BR, Dekker J, Mirny LA (2012) Iterative correction of Hi-C data reveals hallmarks of chromosome organization. Nat Methods, 9: 999–1003

34. Janissen R, Barth R, Polinder M, van der Torre J, Dekker C (2024) Single-molecule visualization of twin-supercoiled domains generated during transcription. Nucleic Acids Res, 52: 1677–1687

35. Jeppsson K, Pradhan B, Sutani T, Sakata T, Umeda Igarashi M, Berta DG, Kanno T, Nakato R, Shirahige K, Kim E, Björkegren C (2024) Loop-extruding Smc5/6 organizes transcription-induced positive DNA supercoils. Mol Cell, 84: 867–882.e5

36. Jupe ER, Sinden RR, Cartwright IL (1993) Stably maintained microdomain of localized unrestrained supercoiling at a Drosophila heat shock gene locus. EMBO J, 12: 1067–1075

37. Kagey MH, Newman JJ, Bilodeau S, Zhan Y, Orlando DA, van Berkum NL, Ebmeier CC, Goossens J, Rahl PB, Levine SS, Taatjes DJ, Dekker J, Young RA (2010) Mediator and cohesin connect gene expression and chromatin architecture. Nature, 467: 430–435

38. Kim D, Pertea G, Trapnell C, Pimentel H, Kelley R, Salzberg SL (2013) TopHat2: accurate alignment of transcriptomes in the presence of insertions, deletions and gene fusions. Genome Biol, 14: R36

39. Kim SH, Ganji M, Kim E, van der Torre J, Abbondanzieri E, Dekker C (2018) DNA sequence encodes the position of DNA supercoils. Elife, 7: e36557

40. Kim TK, Hemberg M, Gray JM, Costa AM, Bear DM, Wu J, Harmin DA, Laptewicz M, Barbara-Haley K, Kuersten S, Markenscoff-Papadimitriou E, Kuhl D, Bito H, Worley PF, Kreiman G, Greenberg ME (2010) Widespread transcription at neuronal activity-regulated enhancers. Nature, 465: 182–187

41. Koch F, Fenouil R, Gut M, Cauchy P, Albert TK, Zacarias-Cabeza J, Spicuglia S, de la Chapelle AL, Heidemann M, Hintermair C, Eick D, Gut I, Ferrier P, Andrau J-C (2011) Transcription initiation platforms and GTF recruitment at tissue-specific enhancers and promoters. Nat Struct Mol Biol, 18: 956–963

42. Kouzine F, Gupta A, Baranello L, Wojtowicz D, Ben-Aissa K, Liu J, Przytycka TM, Levens D (2013) Transcription-dependent dynamic supercoiling is a short-range genomic force. Nat Struct Mol Biol, 20: 396–403

43. Kouzine F, Liu J, Sanford S, Chung HJ, Levens D (2004) The dynamic response of upstream DNA to transcription-generated torsional stress. Nat Struct Mol Biol, 11: 1092–1100

44. Kouzine F, Sanford S, Elisha-Feil Z, Levens D (2008) The functional response of upstream DNA to dynamic supercoiling in vivo. Nat Struct Mol Biol, 15: 146–154

45. Krassovsky K, Ghosh RP, Meyer BJ (2021) Genome-wide profiling reveals functional interplay of DNA sequence composition, transcriptional activity, and nucleosome positioning in driving DNA supercoiling and helix destabilization in C. elegans. Genome Res, 31: 1187–1202

46. Langmead B, Salzberg SL (2012) Fast gapped-read alignment with Bowtie 2. Nat Methods, 9: 357–359

47. Lawrence M, Gentleman R, Carey V (2009) rtracklayer: an R package for interfacing with genome browsers. Bioinformatics, 25: 1841–1842

48. Lawrence M, Huber W, Pagès H, Aboyoun P, Carlson M, Gentleman R, Morgan MT, Carey VJ (2013) Software for computing and annotating genomic ranges. PLoS Comput Biol, 9: e1003118

49. Lettice LA, Heaney SJ, Purdie LA, Li L, de Beer P, Oostra BA, Goode D, Elgar G, Hill RE, de Graaff E (2003) A long-range Shh enhancer regulates expression in the developing limb and fin and is associated with preaxial polydactyly. Hum Mol Genet, 12: 1725–1735

50. Li H, Handsaker B, Wysoker A, Fennell T, Ruan J, Homer N, Marth G, Abecasis G, Durbin R, 1000 GPDPS (2009) The Sequence Alignment/Map format and SAMtools. Bioinformatics, 25: 2078–2079

51. Li H (2013) Aligning sequence reads, clone sequences and assembly contigs with BWA-MEM. arXiv, 1303.3997v2

52. Lieberman-Aiden E, van Berkum NL, Williams L, Imakaev M, Ragoczy T, Telling A, Amit I, Lajoie BR, Sabo PJ, Dorschner MO, Sandstrom R, Bernstein B, Bender MA, Groudine M, Gnirke A, Stamatoyannopoulos J, Mirny LA, Lander ES, Dekker J (2009) Comprehensive mapping of long-range interactions reveals folding principles of the human genome. Science, 326: 289–293

53. Liu LF, Wang JC (1987) Supercoiling of the DNA template during transcription. Proc Natl Acad Sci U S A, 84: 7024–7027

54. Longo GMC, Sayols S, Stefanova ME, Xie T, Elsayed W, Panagi A, Stavridou AI, Petrosino G, Ing-Simmons E, Melo US, Gothe HJ, Vaquerizas JM, Kotini AG, Papantonis A, Mundlos S, Roukos V (2024) Type II topoisomerases shape multi-scale 3D chromatin folding in regions of positive supercoils. Mol Cell, 84: 4267–4281.e8

55. Lupiáñez DG, Kraft K, Heinrich V, Krawitz P, Brancati F, Klopocki E, Horn D, Kayserili H, Opitz JM, Laxova R, Santos-Simarro F, Gilbert-Dussardier B, Wittler L, Borschiwer M, Haas SA, Osterwalder M, Franke M, Timmermann B, Hecht J, Spielmann M, Visel A, Mundlos S (2015) Disruptions of topological chromatin domains cause pathogenic rewiring of gene-enhancer interactions. Cell, 161: 1012–1025

56. Lyu X, Rowley MJ, Corces VG (2018) Architectural Proteins and Pluripotency Factors Cooperate to Orchestrate the Transcriptional Response of hESCs to Temperature Stress. Mol Cell, 71: 940–955.e7

57. Ma J, Bai L, Wang MD (2013) Transcription under torsion. Science, 340: 1580–1583

58. Mattingly M, Seidel C, Muñoz S, Hao Y, Zhang Y, Wen Z, Florens L, Uhlmann F, Gerton JL (2022) Mediator recruits the cohesin loader Scc2 to RNA Pol II-transcribed genes and promotes sister chromatid cohesion. Curr Biol, 32: 2884–2896.e6

59. Meuleman W, Peric-Hupkes D, Kind J, Beaudry JB, Pagie L, Kellis M, Reinders M, Wessels L, van Steensel B (2013) Constitutive nuclear lamina-genome interactions are highly conserved and associated with A/T-rich sequence. Genome Res, 23: 270–280

60. Naughton C, Avlonitis N, Corless S, Prendergast JG, Mati IK, Eijk PP, Cockroft SL, Bradley M, Ylstra B, Gilbert N (2013) Transcription forms and remodels supercoiling domains unfolding large-scale chromatin structures. Nat Struct Mol Biol, 20: 387–395

61. Neguembor MV, Martin L, Castells-García Á, Gómez-García PA, Vicario C, Carnevali D, AlHaj Abed J, Granados A, Sebastian-Perez R, Sottile F, Solon J, Wu CT, Lakadamyali M, Cosma MP (2021) Transcription-mediated supercoiling regulates genome folding and loop formation. Mol Cell, 81: 3065–3081.e12

62. Niwa H, Miyazaki J, Smith AG (2000) Quantitative expression of Oct-3/4 defines differentiation, dedifferentiation or self-renewal of ES cells. Nat Genet, 24: 372–376

63. Nora EP, Lajoie BR, Schulz EG, Giorgetti L, Okamoto I, Servant N, Piolot T, van Berkum NL, Meisig J, Sedat J, Gribnau J, Barillot E, Blüthgen N, Dekker J, Heard E (2012) Spatial partitioning of the regulatory landscape of the X-inactivation centre. Nature, 485: 381–385

64. O’Sullivan JM, Tan-Wong SM, Morillon A, Lee B, Coles J, Mellor J, Proudfoot NJ (2004) Gene loops juxtapose promoters and terminators in yeast. Nat Genet, 36: 1014–1018

65. Phillips-Cremins JE, Sauria ME, Sanyal A, Gerasimova TI, Lajoie BR, Bell JS, Ong CT, Hookway TA, Guo C, Sun Y, Bland MJ, Wagstaff W, Dalton S, McDevitt TC, Sen R, Dekker J, Taylor J, Corces VG (2013) Architectural protein subclasses shape 3D organization of genomes during lineage commitment. Cell, 153: 1281–1295

66. Pickersgill H, Kalverda B, de Wit E, Talhout W, Fornerod M, van Steensel B (2006) Characterization of the Drosophila melanogaster genome at the nuclear lamina. Nat Genet, 38: 1005–1014

67. Pommier Y, Nussenzweig A, Takeda S, Austin C (2022) Human topoisomerases and their roles in genome stability and organization. Nat Rev Mol Cell Biol, 23: 407–427

68. Pommier Y, Sun Y, Huang SN, Nitiss JL (2016) Roles of eukaryotic topoisomerases in transcription, replication and genomic stability. Nat Rev Mol Cell Biol, 17: 703–721

69. Pyne ALB, Noy A, Main KHS, Velasco-Berrelleza V, Piperakis MM, Mitchenall LA, Cugliandolo FM, Beton JG, Stevenson CEM, Hoogenboom BW, Bates AD, Maxwell A, Harris SA (2021) Base-pair resolution analysis of the effect of supercoiling on DNA flexibility and major groove recognition by triplex-forming oligonucleotides. Nat Commun, 12: 1053

70. Racko D, Benedetti F, Dorier J, Stasiak A (2018) Transcription-induced supercoiling as the driving force of chromatin loop extrusion during formation of TADs in interphase chromosomes. Nucleic Acids Res, 46: 1648–1660

71. Racko D, Benedetti F, Dorier J, Stasiak A (2019) Are TADs supercoiled? Nucleic Acids Res, 47: 521–532

72. Ramasamy S, Aljahani A, Karpinska MA, Cao TBN, Velychko T, Cruz JN, Lidschreiber M, Oudelaar AM (2023) The Mediator complex regulates enhancer-promoter interactions. Nat Struct Mol Biol, 30: 991–1000

73. Ramírez F, Bhardwaj V, Arrigoni L, Lam KC, Grüning BA, Villaveces J, Habermann B, Akhtar A, Manke T (2018) High-resolution TADs reveal DNA sequences underlying genome organization in flies. Nat Commun, 9: 189

74. Ramírez F, Ryan DP, Grüning B, Bhardwaj V, Kilpert F, Richter AS, Heyne S, Dündar F, Manke T (2016) deepTools2: a next generation web server for deep-sequencing data analysis. Nucleic Acids Res, 44: W160–5

75. Rao SS, Huntley MH, Durand NC, Stamenova EK, Bochkov ID, Robinson JT, Sanborn AL, Machol I, Omer AD, Lander ES, Aiden EL (2014) A 3D map of the human genome at kilobase resolution reveals principles of chromatin looping. Cell, 159: 1665–1680

76. Reimand J, Kull M, Peterson H, Hansen J, Vilo J (2007) g:Profiler--a web-based toolset for functional profiling of gene lists from large-scale experiments. Nucleic Acids Res, 35: W193–200

77. Robinson MD, McCarthy DJ, Smyth GK (2010) edgeR: a Bioconductor package for differential expression analysis of digital gene expression data. Bioinformatics, 26: 139–140

78. Robinson MD, Oshlack A (2010) A scaling normalization method for differential expression analysis of RNA-seq data. Genome Biol, 11: R25

79. Rowley MJ, Lyu X, Rana V, Ando-Kuri M, Karns R, Bosco G, Corces VG (2019) Condensin II Counteracts Cohesin and RNA Polymerase II in the Establishment of 3D Chromatin Organization. Cell Rep, 26: 2890–2903.e3

80. Rowley MJ, Nichols MH, Lyu X, Ando-Kuri M, Rivera ISM, Hermetz K, Wang P, Ruan Y, Corces VG (2017) Evolutionarily Conserved Principles Predict 3D Chromatin Organization. Mol Cell, 67: 837–852.e7

81. Saffran WA, Welsh JT, Knobler RM, Gasparro FP, Cantor CR, Edelson RL (1988) Preparation and characterization of biotinylated psoralen. Nucleic Acids Res, 16: 7221–7231

82. Sexton T, Yaffe E, Kenigsberg E, Bantignies F, Leblanc B, Hoichman M, Parrinello H, Tanay A, Cavalli G (2012) Three-dimensional folding and functional organization principles of the Drosophila genome. Cell, 148: 458–472

83. Sinden RR, Bat O, Kramer PR (1999) Psoralen cross-linking as probe of torsional tension and topological domain size in vivo. Methods, 17: 112–124

84. Sinden RR, Carlson JO, Pettijohn DE (1980) Torsional tension in the DNA double helix measured with trimethylpsoralen in living E. coli cells: analogous measurements in insect and human cells. Cell, 21: 773–783

85. Smith CL, Kubo M, Imamoto F (1978) Promoter-specific inhibition of transcription by antibiotics which act on DNA gyrase. Nature, 275: 420–423

86. Spicuglia S, Vanhille L (2012) Chromatin signatures of active enhancers. Nucleus, 3: 126–131

87. Tan-Wong SM, Zaugg JB, Camblong J, Xu Z, Zhang DW, Mischo HE, Ansari AZ, Luscombe NM, Steinmetz LM, Proudfoot NJ (2012) Gene loops enhance transcriptional directionality. Science, 338: 671–675

88. Tedeschi A, Wutz G, Huet S, Jaritz M, Wuensche A, Schirghuber E, Davidson IF, Tang W, Cisneros DA, Bhaskara V, Nishiyama T, Vaziri A, Wutz A, Ellenberg J, Peters JM (2013) Wapl is an essential regulator of chromatin structure and chromosome segregation. Nature, 501: 564–568

89. Teo AK, Arnold SJ, Trotter MW, Brown S, Ang LT, Chng Z, Robertson EJ, Dunn NR, Vallier L (2011) Pluripotency factors regulate definitive endoderm specification through eomesodermin. Genes Dev, 25: 238–250

90. Teves SS, Henikoff S (2014) Transcription-generated torsional stress destabilizes nucleosomes. Nat Struct Mol Biol, 21: 88–94

91. Uusküla-Reimand L, Hou H, Samavarchi-Tehrani P, Rudan MV, Liang M, Medina-Rivera A, Mohammed H, Schmidt D, Schwalie P, Young EJ, Reimand J, Hadjur S, Gingras A-C, Wilson MD (2016) Topoisomerase II beta interacts with cohesin and CTCF at topological domain borders. Genome Biology, 17:

92. Vallier L, Touboul T, Brown S, Cho C, Bilican B, Alexander M, Cedervall J, Chandran S, Ahrlund-Richter L, Weber A, Pedersen RA (2009) Signaling pathways controlling pluripotency and early cell fate decisions of human induced pluripotent stem cells. Stem Cells, 27: 2655–2666

93. Valton AL, Venev SV, Mair B, Khokhar ES, Tong AHY, Usaj M, Chan K, Pai AA, Moffat J, Dekker J (2022) A cohesin traffic pattern genetically linked to gene regulation. Nat Struct Mol Biol, 29: 1239–1251

94. Vietri Rudan M, Barrington C, Henderson S, Ernst C, Odom DT, Tanay A, Hadjur S (2015) Comparative Hi-C reveals that CTCF underlies evolution of chromosomal domain architecture. Cell Rep, 10: 1297–1309

95. Visser BJ, Sharma S, Chen PJ, McMullin AB, Bates ML, Bates D (2022) Psoralen mapping reveals a bacterial genome supercoiling landscape dominated by transcription. Nucleic Acids Research, 50: 4436–4449

96. Whyte WA, Orlando DA, Hnisz D, Abraham BJ, Lin CY, Kagey MH, Rahl PB, Lee TI, Young RA (2013) Master transcription factors and mediator establish super-enhancers at key cell identity genes. Cell, 153: 307–319

97. Yao Q, Zhu L, Shi Z, Banerjee S, Chen C (2025) Topoisomerase-modulated genome-wide DNA supercoiling domains colocalize with nuclear compartments and regulate human gene expression. Nat Struct Mol Biol, 32: 48–61

98. Yiangou L, Grandy RA, Morell CM, Tomaz RA, Osnato A, Kadiwala J, Muraro D, Garcia-Bernardo J, Nakanoh S, Bernard WG, Ortmann D, McCarthy DJ, Simonic I, Sinha S, Vallier L (2019) Method to Synchronize Cell Cycle of Human Pluripotent Stem Cells without Affecting Their Fundamental Characteristics. Stem Cell Reports, 12: 165–179

99. Zheng H, Xie W (2019) The role of 3D genome organization in development and cell differentiation. Nat Rev Mol Cell Biol, 20: 535–550

